# High Force Catch Bond Mechanism of Bacterial Adhesion in the Human Gut

**DOI:** 10.1101/2020.01.21.913590

**Authors:** Zhaowei Liu, Haipei Liu, Andrés M. Vera, Rafael C. Bernardi, Philip Tinnefeld, Michael A. Nash

**Affiliations:** Institute of Physical Chemistry, Department of Chemistry, University of Basel, 4058 Basel, Switzerland; Department of Biosystems Science and Engineering, ETH Zurich, 4058 Basel, Switzerland; Faculty of Chemistry and Pharmacy, NanoBioScience, Ludwig-Maximilians-Universität München, Munich, Germany; Beckman Institute for Advanced Science and Technology, University of Illinois at Urbana-Champaign, 61801 Urbana, IL, USA

**Author notes:** **Corresponding Author** (M. A. N).

## Abstract

Bacterial colonization of the human intestine requires firm adhesion of bacteria to insoluble targets under hydrodynamic flow. Here we report the molecular mechanism behind an mechanostable protein complex responsible for resisting high shear forces and adhering bacteria to cellulose fibers in the human gut. Using single-molecule force spectroscopy (SMFS), single-molecule FRET (smFRET), and molecular dynamics (MD) simulations, we resolved two binding modes and three unbinding reaction pathways of a mechanically ultrastable *R. champanellensis* (*Rc*) Dockerin-Cohesin (Doc-Coh) complex. The complex assembles in two discrete binding modes with significantly different mechanical properties, with one breaking at ~500 pN and the other at ~200 pN at loading rates from 1-100 nN/sec. A neighboring X-module domain allosterically regulates the binding interaction and inhibits one of the low-force pathways at high loading rates, giving rise to a new mechanism of catch bonding that manifests under force ramp protocols. Multi-state Monte Carlo simulations show strong agreement with experimental results, validating the proposed kinetic scheme. These results explain mechanistically how gut microbes regulate cell adhesion strength at high shear stress through intricate molecular mechanisms including dual-binding modes, mechanical allostery and catch bonds.

---

When cells adhere to surfaces under flow, adhesion bonds at the cell-surface interface experience mechanical tension and resist hydrodynamic drag forces. Because of this mechanical selection pressure, adhesion proteins have evolved molecular mechanisms to deal with tension in different ways. Most bonds not involved in force transduction *in vivo* have lifetimes that decay exponentially with applied force, a behavior well described by the classical Bell-Evans slip bond model^1–3^. Less intuitive are catch bonds^4–7^, which are receptor-ligand interactions that serve as band pass filters for force perturbations, becoming stronger with applied force and weakening when force is released. When probed in constant force mode, the lifetime of a catch bond will rise as the force setpoint is increased. When probed in force ramp mode or constant speed mode, catch bonds typically give rise to bimodal rupture force distributions^8,9^. Different kinetic state models and network topologies can be used to describe catch bonds^10^. For example, mechanical allostery models such as the one-state two-pathway model^11^, or two independent sites model^18,12^ have been applied to mathematically describe catch bond behavior^6,8,12,13^.

The *R. champanellensis (Rc)* cellulosome^14,15^ is a bacterial protein complex found in the human gut that adheres to and digests plant fiber. The large supramolecular complex is held together by Dockerin-Cohesin (Doc:Coh) interactions^16^ which comprise a family of homologous high-affinity receptor-ligand pairs. A limited number of Doc:Coh complexes are known to exhibit dual-binding modes^17–21^ where the complex populates two distinct binding conformations involving different sets of binding residues on Doc recognizing the same residues on Coh. The two binding modes have nearly identical equilibrium binding affinity, making them challenging to observe experimentally using conventional bulk experiments. Instead, we took a single-molecule approach which is uniquely suited for studying such discrete heterogeneous systems. Single-molecule force spectroscopy with the atomic force microscope (AFM-SMFS) is able to explore a large force range up to several nN and has been used to characterize protein folding pathways^22–24^ and receptor-ligand interactions^25–28^. Single-molecule FRET (smFRET) is capable of measuring distances at the molecular scale^29^, and has been used to study protein dynamics^30,31^ and to characterize structures of receptor-ligand complexes^32,33^. At the computational level, these experimental single-molecule approaches can be elaborated upon by employing molecular dynamics (MD) simulations^34^. When combined, these experimental and computational approaches can provide mechanistic insights into the dynamics of receptor-ligand complexes^26,35^.

Here we investigated a complex called XMod-Doc:Coh responsible for anchoring the *Rc* cellulosome complex to the bacterial cell wall and used AFM-SMFS, smFRET, and MD simulations to study putative dual-binding modes and catch bond behavior of the complex. We developed a state map with experimentally transition rates to fully describe the system, and performed kinetic Monte Carlo simulations that recapitulate the experimental data. What emerges from this three state kinetic scheme is a picture of a unique adhesion bond that resembles a catch bond when probed under force ramp conditions, but maintains slip bonding under constant force.

## Results

### XMod-Doc:Coh homology model and expression cassettes

A proposed topology of the *Rc* cellulosome is shown in Fig. 1a^14,15^. We investigated the interaction between Dockerin B (Doc) located at the C-terminus of Scaffoldin B, and Cohesin E (Coh) which is covalently attached to the peptidoglycan cell wall. Adjacent to Doc is an Ig-like domain called X-module (XMod; Fig. 1a, purple). This newly reported complex is homologous to previously reported complexes from *R. flavefaciens* (*Rf*)^35–37^. Full amino acid sequences are given in the Supplementary Information. Since no structural information was available for the *Rc* XMod-Doc:Coh complex, using Modeller 9.22^38^ we created homology models of each protein domain. The structure of the *Rc* XMod-Doc domain was modeled based on the available structure of *Rf* CttA XMod-Doc (PDB 4IU3)^39^, which shares a 20% sequence identity (35% similarity) with the *Rc* domain. The structure of the *Rc* Coh domain was modeled based on two different available structures from *Rf*, namely CohE (PDB 4IU3) with 15% sequence identity (28% similarity), and CohG (PDB 4WKZ)^40^ with 18% sequence identity (34% similarity). The 10 models with highest score from Modeller were selected for each domain/template pair, resulting in 10 models for the *Rc* XMod-Doc domain, and 20 models for the *Rc* Coh domain. Employing VMD^41^, we assembled 200 models of the *Rc* XMod-Doc:Coh complex in each of the two binding modes (Fig. 1b and c), building all possible combinations between XMod-Doc and Coh models. For binding mode A, the structure of the *Rf* XMod-Doc:Coh complex (PDB 4IU3) was employed to guide the *Rc* Coh:Doc interface alignment. To create a model for the hypothesized alternative binding mode B, Doc helix 1 from the homology model structure was used as a guide for the superposition of Doc helix 3. This alignment resulted in the XMod-Doc rotating 180° with respect to Coh. The models show that Doc binds Coh via the two Ca^2+^ binding loops and two binding helices (helices 1 and 3, see **Fig. S1**), which are connected by a short helix 2. This duplicated F-hand motif is consistent with those of other Doc domains which have been shown to exhibit dual binding modes^17,19–21^.

**Figure 1.**
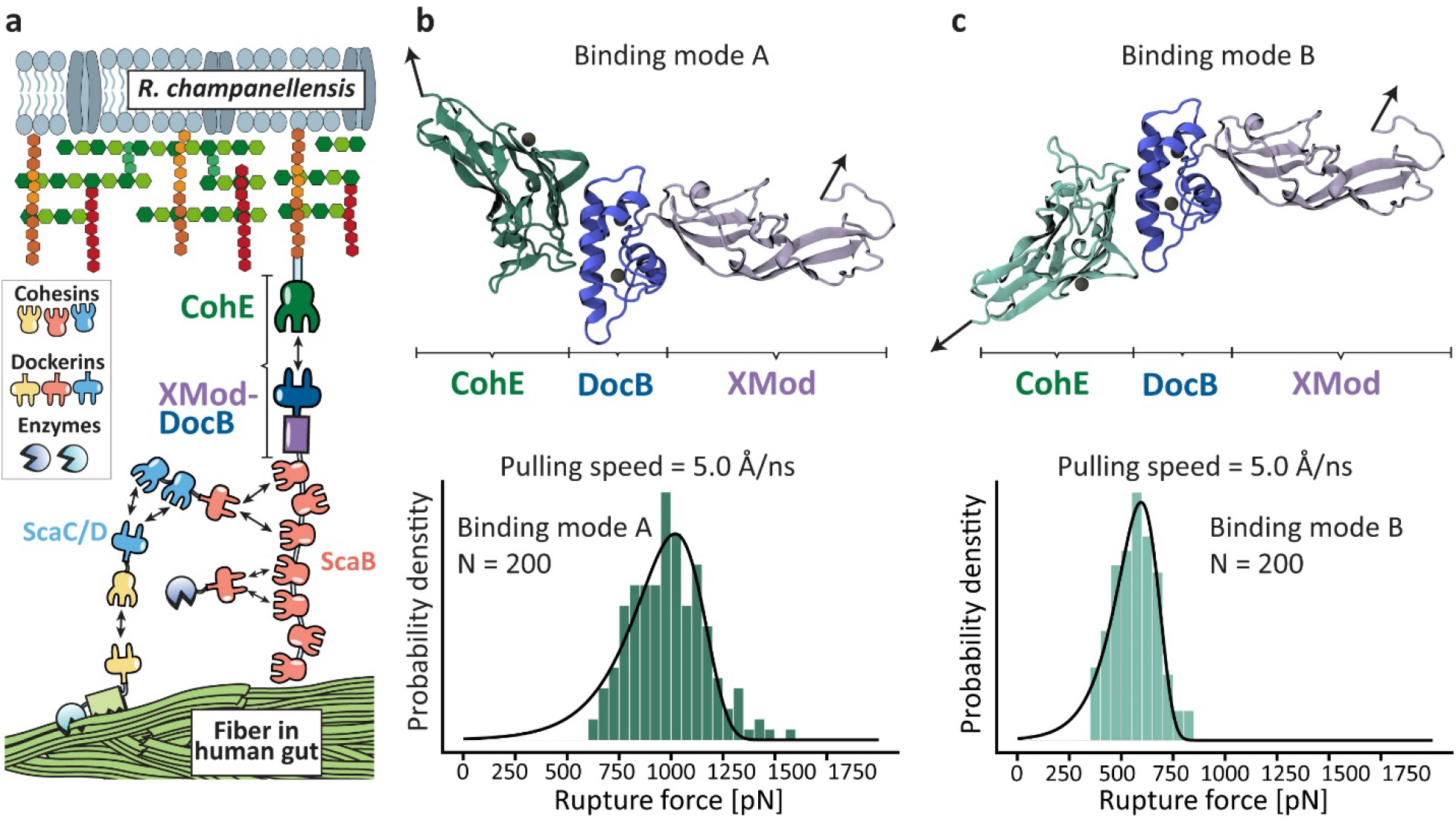
*Rc* XMod-Doc:Coh complex, dual binding modes, and molecular dynamics simulation of complex dissociation. **a:** The *Rc*-cellulosomal network is assembled through interactions between Doc and Coh domains. Cellulose binding domains, digestive enzymes, and structural scaffold proteins self-assemble into a cellulosome complex, which binds and digests cellulose fibers in the human gut. XMod-DocB and CohE form a mechanically stable protein complex that anchors the cellulosomal network to the cell surface. **b and c:** Structural models showing the XMod-Doc:Coh complex in the two hypothesized binding modes. Green: Coh, blue: Doc, purple: XMod. Calcium ions are shown as black spheres. In both binding modes, the rupture forces observed for the 5 most stable models were measured by performing 200 steered molecular dynamics (SMD) replicas and plotting as histograms. Rupture force histograms were fitted with the Bell-Evans distribution. The most probable rupture force was 1020 pN in binding mode A (panel **b**) and 595 pN in binding mode B (panel **c**) at a pulling speed of 5.0 Å/ns.

We cloned polyproteins containing several modules for AFM-SMFS and purified them from *E. coli*. A ddFLN4 and an elastin-like polypeptide (ELP) were used as an unfolding fingerprint domain^42^ and flexible linker^43–46,^ respectively. The Coh construct (N-to C-terminus) was Coh-ddFLN4-ELP-HIS-ybbr. The XMod-Doc construct (N-to C-terminus) was ybbr-ELP-ddFLN4-XMod-Doc-HIS. The ybbR tag facilitated site-specific and covalent linkage to the coverglass or cantilever tip^47^. The loading geometry with Coh pulled from its C-terminus and XMod-Doc pulled from its N-terminus precisely mimicked that experienced by the complex *in vivo*. Analysis of the equilibrium binding affinity of WT XMod-Doc:Coh using isothermal titration calorimetry (ITC) revealed *K*_*D*_=1.0 ± 0.3 nM and a binding stoichiometry of 1:1. SDS-PAGE and mass spectrometry analysis indicated a molecular weight of 44 kD for Coh construct and 55 kD for XMod-Doc construct.

### Steered molecular dynamics simulations reveals a weak and a strong binding mode

To examine the stability of *Rc* XMod-Doc:Coh under mechanical load we carried out steered molecular dynamics (SMD) simulations^3^ employing NAMD^48,49^ and its QwikMD^50^ interface. First, to test the stability of the 200 models of the complex in each binding mode, we performed equilibrium MD simulations for a combined simulation time of 2.0 μs, followed by a combined 8.0 μs of SMD simulations at constant pulling velocity. These SMD simulations served as a metric to eliminate unsuitable structural models. We expected that good structural models should be stable under mechanical load, therefore, for each binding mode, we selected the 5 strongest complexes out of the 200 models. In fact, some of the 400 complexes were found not to be stable already after the equilibrium MD, and due to the low sequence identity of the templates, most of the models were not stable under mechanical load. A visual observation in VMD showed that many of these models had only partial contact between Coh and Doc following equilibrium MD. From the 5 strongest models for each binding mode, we performed 200 production SMD simulation replicas, using a similar protocol as previously described^26,35^. The simulations reveal that the dissociation of XMod-Doc:Coh occurs at clearly distinct forces for the two different binding modes, with mode A dissociating at ~1020 pN, and mode B at ~595 pN, both at a 5.0 Å/ns pulling speed (Fig. 1b and c).

### Wild type XMod-Doc:Coh unbinds along 3 distinct pathways

We performed AFM-SMFS with Coh covalently attached to the cantilever tip through its C-terminal ybbR tag and XMod-Doc covalently attached to the surface (Fig. 2a) through its N-terminal ybbR tag. The XMod-Doc:Coh complex was formed by approaching the AFM tip to the surface and dwelling for 200 ms. After XMod-Doc:Coh complex formation, the cantilever base was retracted at constant speed and a force-extension curve was recorded. This procedure was repeated thousands of times typically over a 12 hours period to generate large datasets of force *vs.* extension curves. The recorded force curves were transformed into force vs. contour length space using a freely rotating chain (FRC) elasticity model (Eq. S1). We searched for the contour length pattern of ddFLN4, which contained ~32 nm of total contour length that resulted from a two-step unfolding pattern. Since one ddFLN4 molecule was contained in the surface-linked protein, and another one in the cantilever-linked protein, we only analyzed curves which contained in total two ddFLN4 unfolding fingerprints, thereby eliminating spurious signals.

**Figure 2.**
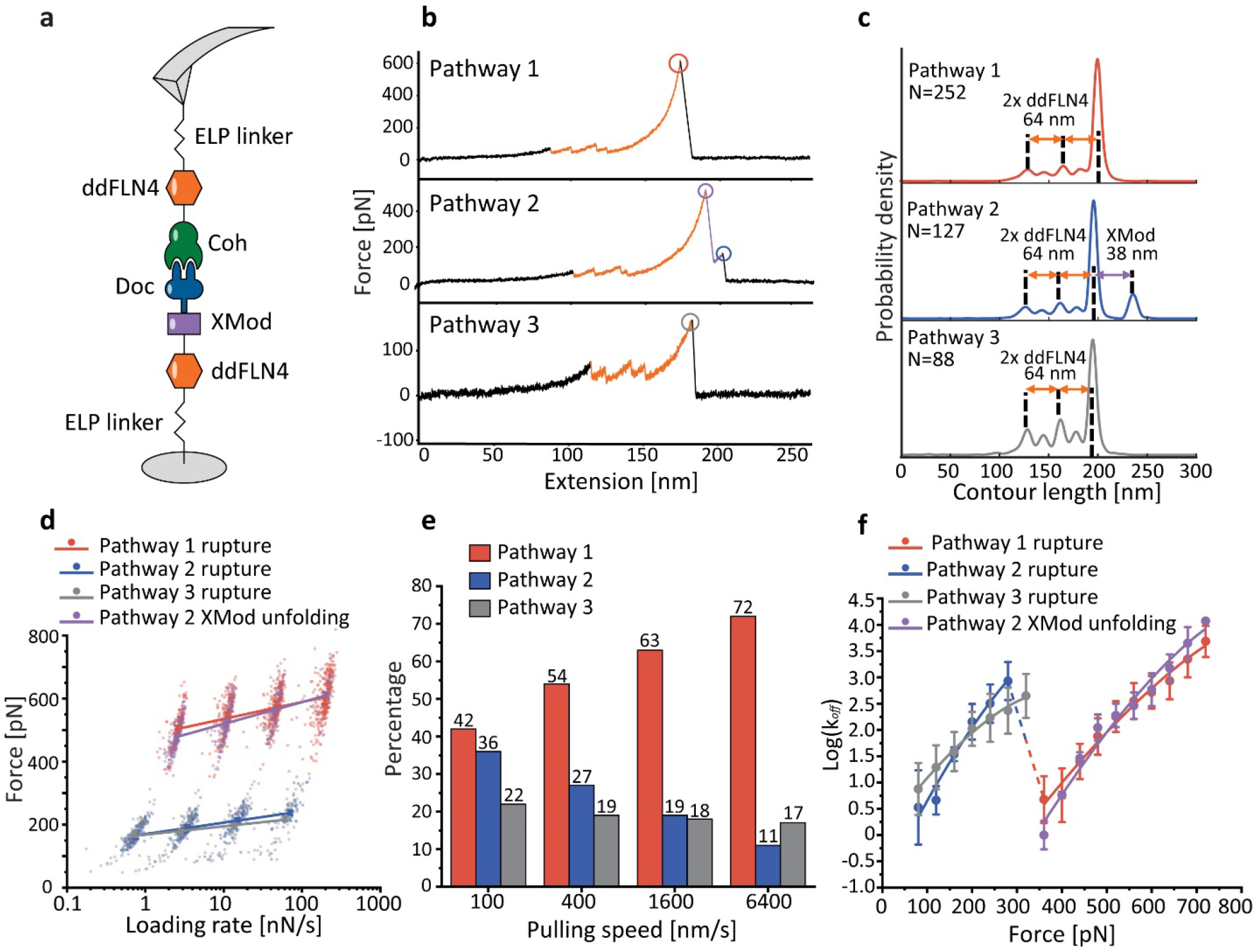
XMod-Doc:Coh unbinds along three pathways under mechanical load. **a:** Experimental configuration with Coh-ddFLN4-ELP immobilized on the AFM tip and ELP-ddFLN4-XMod-Doc immobilized on the surface. Immobilization was site-specific and covalent through a terminal ybbR tag on the ELP. **b:** Three different classes of force curves were repeatedly observed, corresponding to different pathways. In pathway 1 (P1), the complex ruptured at high force (500-600 pN, red) with the XMod remaining folded. In pathway 2 (P2), XMod unfolded (purple) followed by a low force rupture of the complex (blue). In pathway 3 (P3), the complex ruptured at low force rupture (grey) with the XMod remaining folded. Unfolding of the two ddFLN4 fingerprint domains (orange) was used to identify single-molecule traces. **c:** Combined contour length histograms for each unbinding pathway. Force-extension traces were transformed using a freely rotating chain elasticity model and aligned using cross-correlation analysis. Histograms show contour length increments resulting from unfolding of 2x ddFLN4 and XMod. **d:** Rupture force vs. loading rate plot showing final XMod-Doc:Coh complex rupture events obtained from the three pathways, as well as XMod unfolding events observed in P2. Lines show linear Bell-Evans fits of the most probable rupture/unfolding force vs. logarithm of loading rate to obtain *Δx*^*‡*^ and *k0* (Eq. S2). Fitted *Δx*^*‡*^ and *k0* values are listed in Table 1. **e:** Percentages of the three pathways in all rupture events at different pulling speeds. **f:** Kinetic off-rate (*koff*) *vs.* force for complex rupture events. Off-rates were calculated using the histogram transformation method (Eq. S3). Lines show fitting to the analytical expression (Eq. S5). The fitted *Δx*^*‡*^, *ΔG*^*‡*^ and *k0* values are listed in Table 1.

We repeatedly observed three distinct unbinding pathways of the complex, as shown in Fig. 2b. We refer to these as pathway 1 (P1), pathway 2 (P2), and pathway 3 (P3). We used cross-correlation analysis^51,52^ to assemble superposition contour length histograms for each pathway (Fig. 2c). These histograms all showed the distinct unfolding pattern of two ddFLN4 fingerprint domains, adding in total 64 nm contour length to the system. P2 showed an additional 38 nm length increment which matched the expected value for XMod unfolding (116 XMod amino acids * 0.365 nm/amino acid - 5.3 nm folded length = 37 nm) (Fig. 2c, **middle**). The contour length histograms were broadened by occasional unassigned unfolding events that were observed in all three pathways, which we attributed to partial unfolding of Coh. A representative sampling of these unassigned unfolding events are presented in **Fig. S2**.

**Table 1.** Kinetic parameters extracted from AFM-SMFS

| Pathway | Event | $k_0$ [s<sup>-1</sup>]<br>(Bell-Evans) | $\Delta x^\ddagger$ [nm]<br>(Bell-Evans) | $\Delta G^\ddagger$ [k<sub>B</sub>T]<br>(DHS) | $k_0$ [s<sup>-1</sup>]<br>(DHS) | $\Delta x^\ddagger$ [nm]<br>(DHS) |
| --- | --- | --- | --- | --- | --- | --- |
| 1 | High force rupture<br>(XMod folded) | $4.80 \times 10^{-8}$ | 0.178 | 20.0 | $1.57 \times 10^{-4}$ | 0.132 |
| 2 | Unfolding of XMod<br>(measured) | $9.94 \times 10^{-6}$ | 0.139 | 24.0 | $4.79 \times 10^{-6}$ | 0.168 |
| | Unfolding of XMod<br>(corrected) | $4.53 \times 10^{-5}$ | 0.116 | 26.1 | $5.42 \times 10^{-6}$ | 0.157 |
| | Low force rupture<br>after XMod unfold | $1.24 \times 10^{-3}$ | 0.265 | 12.5 | 0.101 | 0.171 |
| 3 | Low force rupture<br>(XMod intact) | $4.18 \times 10^{-5}$ | 0.359 | 7.45 | 1.08 | 0.113 |

Approximately 80% of curves were assigned to P1 or P2. In P1, unfolding of two ddFLN4 domains in series was followed by dissociation of XMod-Doc:Coh at high forces of ~500 pN (Fig. 2b, **top**). In P2, XMod unfolded at high forces, followed by the Doc-Coh complex rupture at low forces of ~200 pN (Fig. 2b, **middle**). This indicated that XMod unfolding significantly destabilized the interaction between Doc and Coh in P2, giving rise to a shielded complex rupture event. The remaining 20% of curves were classified as P3, where XMod-Doc:Coh ruptured at low force (~200 pN, Fig. 2b, **bottom**) and no XMod unfolding was observed. Based on these classifications, we hypothesized that P1 and P2 resulted from complexes with high mechanical stability, which were able to resist external forces as high as ~500 pN prior to high force complex rupture or XMod unfolding. In the cases where XMod unfolded, the Doc-Coh binding interaction became destabilized and ruptured at low force. P3 meanwhile represented a weaker Doc-Coh complex that ruptured at lower force (~200 pN) even without XMod unfolding. The existence of complexes with different mechanical stabilities was consistent with SMD simulation results (Fig. 1c), which indicated dual-binding modes that rupture at distinct forces.

### Allosteric regulation by XMod gives rise to catch bonding in force ramp mode

AFM measurements on WT XMod-Doc:Coh were carried out at pulling speeds of 100, 400, 1600, and 6400 nm/s, which allowed us to investigate the loading rate dependency of complex rupture and XMod unfolding in the various pathways (Fig. 2d). We used the Bell-Evans model (Eq. S2)^1,2^ to analyze the force-loading rate data and obtain the intrinsic off rate (*k*_*0*_) and the distance to the transition state along the reaction coordinate (*Δx*^*‡*^) for the complex rupture events in each pathway, as well as for XMod unfolding along P2 (Table 1).

As shown in Fig. 2e, the percentage of curves that were classified as P3 was independent of the pulling speed, maintaining a value of 17-22% across the range of speeds tested. This observation was consistent with the hypothesis that P3 belonged to a different binding mode than P1 and P2. Interestingly, the ratio between P1 and P2 was dependent on the pulling speed. The likelihood of P1 increased with increasing pulling speed from 100-6400 nm/s, while the likelihood of observing P2 decreased. This means that the complex preferentially populated the pathway with higher rupture force (P1) when pulled at higher loading rates. This switch from low stability P2 to high stability P1 at increasing loading rates is not to be confused with standard scaling based on Bell-Evans theory, which also predicts higher rupture forces at higher loading rates. This behavior, in contrast, represented a discrete non-linear switching from P2 to P1 with much higher rupture forces. Although P1 and P2 rupture events each individually scale as classical slip bonds as a function of the loading rate, the pathway switching behavior precisely mimics that of a catch bond^4,6,12,53,54^ probed under force ramp conditions. In contrast to other reported catch bonds in the literature which occur at low force (<50 pN), XMod-Doc:Coh is activated at much higher forces (>300 pN).

The explanation for this apparent catch bond behavior under force ramp conditions is evident when looking at the loading rate dependency of XMod unfolding. The loading rate dependency of XMod unfolding is steeper than that of the complex rupture in P1 (Fig. 2d, Table 1). Therefore, at high loading rates, far fewer complexes reach sufficiently high forces to unfold XMod prior to complex rupture, thus prohibiting the system from entering P2. This behavior is unique to this particular XMod-Doc:Coh system and was not observed in other Dockerin-Cohesin systems reported thus far^35,36^.

We note that the experimentally observed values for XMod unfolding are slightly biased by the maximal stability of the receptor-ligand complex^55^. This ceiling effect is magnified at high loading rates (>100 nN/sec) because the XMod unfolding force increases and exceeds the maximal force that the complex can withstand. We corrected the XMod unfolding force distribution to take this biasing effect into account^55^, as shown in **Fig. S3**. Using the Bell-Evans model, we obtained the kinetic parameters of XMod unfolding after bias correction (Table 1). This analysis confirmed what was observed in the rupture force *vs.* loading rate scatter plots, namely that XMod has a steeper loading rate dependency (lower *Δx*^*‡*^) than the high force rupture event in P1, and that these scaling differences give rise to catch bonding in force ramp/constant speed mode.

The rupture forces obtained at different pulling speeds were plotted as histograms (**Fig. S4**). We analyzed the complex rupture and XMod unfolding events as irreversible crossings of the system over a single energy barrier according to the formalism of Dudko *et al.*^56,57^ to obtain the intrinsic barrier crossing rate (*k*_*0*_), barrier height (*ΔG*^*‡*^) and distance to the transition state along the reaction coordinate (*Δx*^*‡*^). The complex rupture force distributions from three pathways as well as the corrected XMod unfolding force distribution were transformed into force-dependent off-rate *k*_*off*_*(F)* using Eq. S3. The off-rates were plotted against force and fitted using Eq. S5 to extract the *k*_*0*_, *ΔG*^*‡*^ and *Δx*^*‡*^ values of the various barrier-crossing events, as shown in Fig. 2f and Table 1. The plot of *k*_*off*_ vs. force showed a cross-over regime from 300-400 pN where the off-rate decreased with increasing force, further demonstrating the catch bond behavior arising from the allosteric regulation by XMod.

### AFM-SMFS evidence of dual-binding modes

We hypothesized that P1 and P2 arose from one binding mode, while P3 arose from an alternative binding mode with lower mechanical stability. To test this, we sought to knock out specific binding modes by mutagenesis (Fig. 3a and **Fig. S1**). Using the structural models, we identified key Doc residues likely to be involved in each respective binding mode (**Fig. S1**), and designed mutations to disrupt electrostatics and hydrogen bonding. The mutant designed to knock out binding mode A contained R191A and L195E mutations, and is referred to as BM^A^-KO. The mutant designed to knock out binding mode B contained R140A and M144E mutations and is referred to as BM^B^-KO. Interactions between BM^A^-KO or BM^B^-KO and Coh were then measured using AFM-SMFS at 400 nm/s. For WT XMod-Doc:Coh, the percentage of P3 curves was typically ~20%. As shown in Fig. 3b and **Fig. S5**, BM^A^-KO resulted in a P3 curve percentage that increased to 31%. We attributed this increase in P3 probability to the destabilization of binding mode A, and slight preferential formation of binding mode B as compared to WT. This result indicated that binding mode B was likely associated with the low force pathway P3. However, the mutations were not able to completely knock out binding mode A.

**Figure 3.**
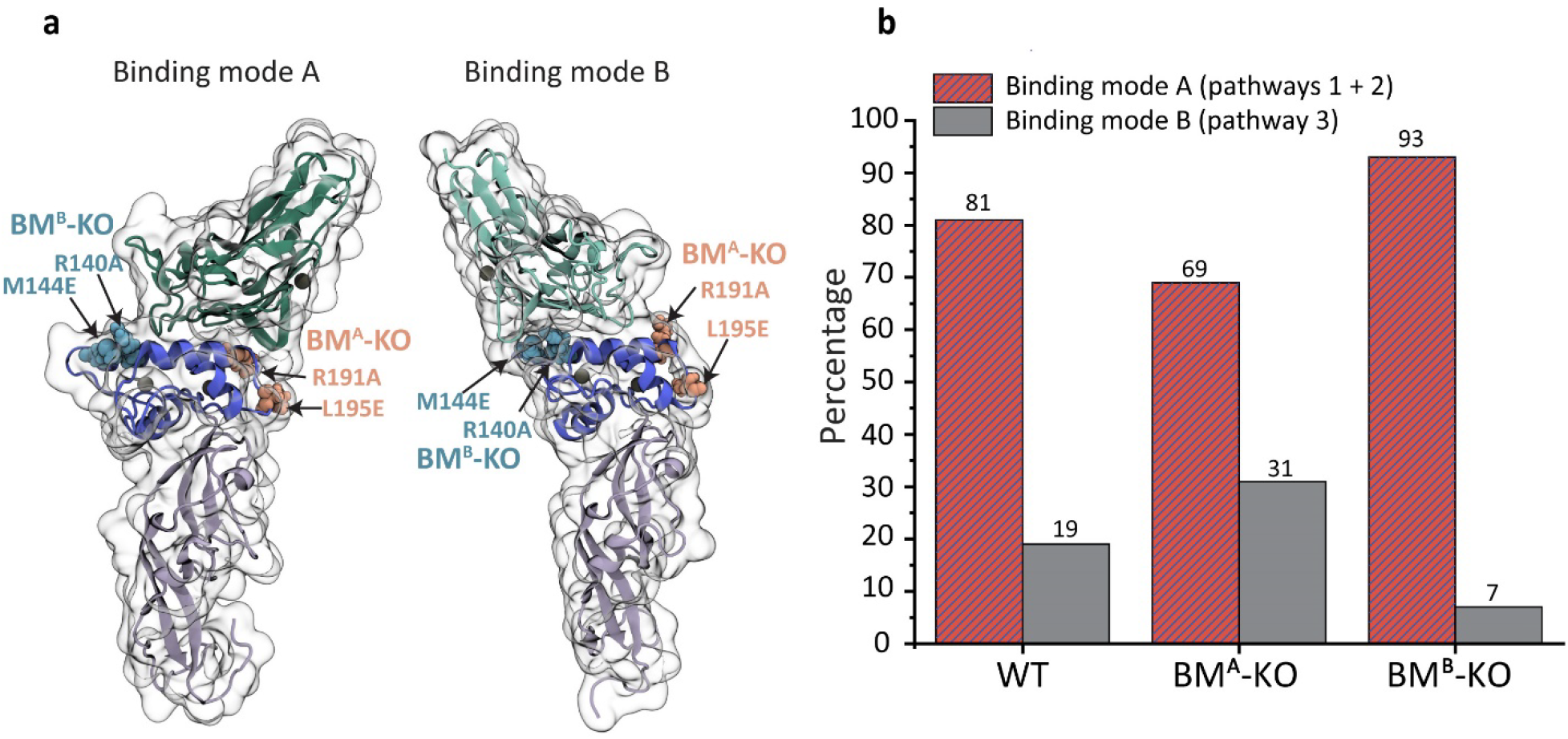
SMFS measurements of XMod-Doc binding mode mutants. **a:** Arginine 191 and leucine 195 on Doc were mutated to alanine and glutamic acid, respectively, to disrupt their interaction with Coh in binding mode A. Arginine 140 and methionine 144 were mutated to alanine and glutamic acid, respectively, to disrupt their interaction with Coh in binding mode B. **b:** Percentages of the two binding modes measured with WT XMod-Doc, BM^A^-KO mutant and BM^B^-KO mutant at 400 nm/s pulling speed. The BM^A^-KO mutant preferentially selects binding mode B while the BM^B^-KO mutant preferentially selects binding mode A.

BM^B^-KO was more effective at knocking out binding activity, and decreased the percentage of P3 curves from 19% for WT down to 7% with a corresponding increase in P1 and P2 percentage. Despite the introduction of destabilizing mutations at the binding interface in BM^B^-KO, we nonetheless obtained a system with higher stability and predominantly high force rupture pathways, a result that may seem counterintuitive but is explained by the presence of a weak binding mode B being knocked out or inhibited by the mutations. Based on these measurements with the binding mode knock-out mutants, we concluded that P1 and P2 are attributable to binding mode A, which is the strong binding mode, while P3 corresponds to binding mode B, which is the weak binding mode.

This conclusion was further supported by a statistical analysis involving a biasing effect of an additional fingerprint domain^55^ (see supplementary information note 2 and **Fig. S6**). We introduced an additional fingerprint domain (I27) whose unfolding force sits in between the P1 and P3 rupture events. If the multi-pathway dissociation behavior that we observed resulted from multiple unbinding reaction pathways from a single bound conformation, we would expect that the likelihood of observing an I27 event would be decorrelated from the pathway classification of the curve. We did not observe this, and instead the vast majority of curves that showed I27 unfolding terminated in a high force rupture event (P1) or XMod unfolding followed by low force rupture (P2). This indicated that complexes that ruptured in a low force rupture event (P3) were not sufficiently strong to unfold I27, consistent with P3 emerging from a discrete binding mode that was weaker than the P1 or P2 complexes, further substantiating the dual binding modes.

### Single-molecule FRET evidence of dual-binding modes

Based on differences in inter-residue distances in the two binding conformations, we used smFRET to observe the dual-binding modes. We introduced a point cysteine mutation at position 154 of Coh and covalently attached a FRET donor dye maleimide-Cy3b. Since XMod-Doc has native cysteines, we used amber suppression^58^ to introduce a non-canonical azide at position 199 of XMod-Doc, and covalently attached DBCO-AF647. Based on the homology models (Fig. 4a), the donor-acceptor distance is expected to be ~3.5 nm in binding mode A and ~4.9 nm in binding mode B. XMod-Doc:Coh complexes were formed by mixing labeled XMod-Doc and Coh in a 1:1 molar ratio and diluting them to ~200 pM. FRET efficiency of individual XMod-Doc:Coh complexes was measured on a confocal microscope and plotted into histograms (Fig. 4b). A bimodal distribution was clearly observed in the FRET efficiency histogram of WT XMod-DocB:Coh, with mean FRET efficiencies of 0.34 and 0.71, corresponding to binding modes B and A, respectively. In addition to labeling and analyzing WT, we introduced the FRET acceptor dye into BM^A^-KO and BM^B^-KO mutants at position 199, and again measured FRET efficiency in complex with labeled Coh using the same protocol as for WT. We found that only the low FRET efficiency peak was observed in BM^A^-KO, meaning that binding mode A corresponding to the high FRET efficiency peak was eliminated by the mutations. The FRET efficiency histogram of BM^B^-KO complexed with Coh meanwhile showed only the high FRET efficiency population, consistent with binding mode B being knocked out. The BM^A^-KO mutant knocks out the binding mode A much more efficiently in smFRET measurement compared to SMFS. We attributed this difference to the acceptor dye destabilizing the complex in binding mode A but not binding mode B, which was confirmed by AFM-SMFS measurements between dye-labeled BM^A^-KO and unlabeled Coh (**Fig. S7**).

**Figure 4.**
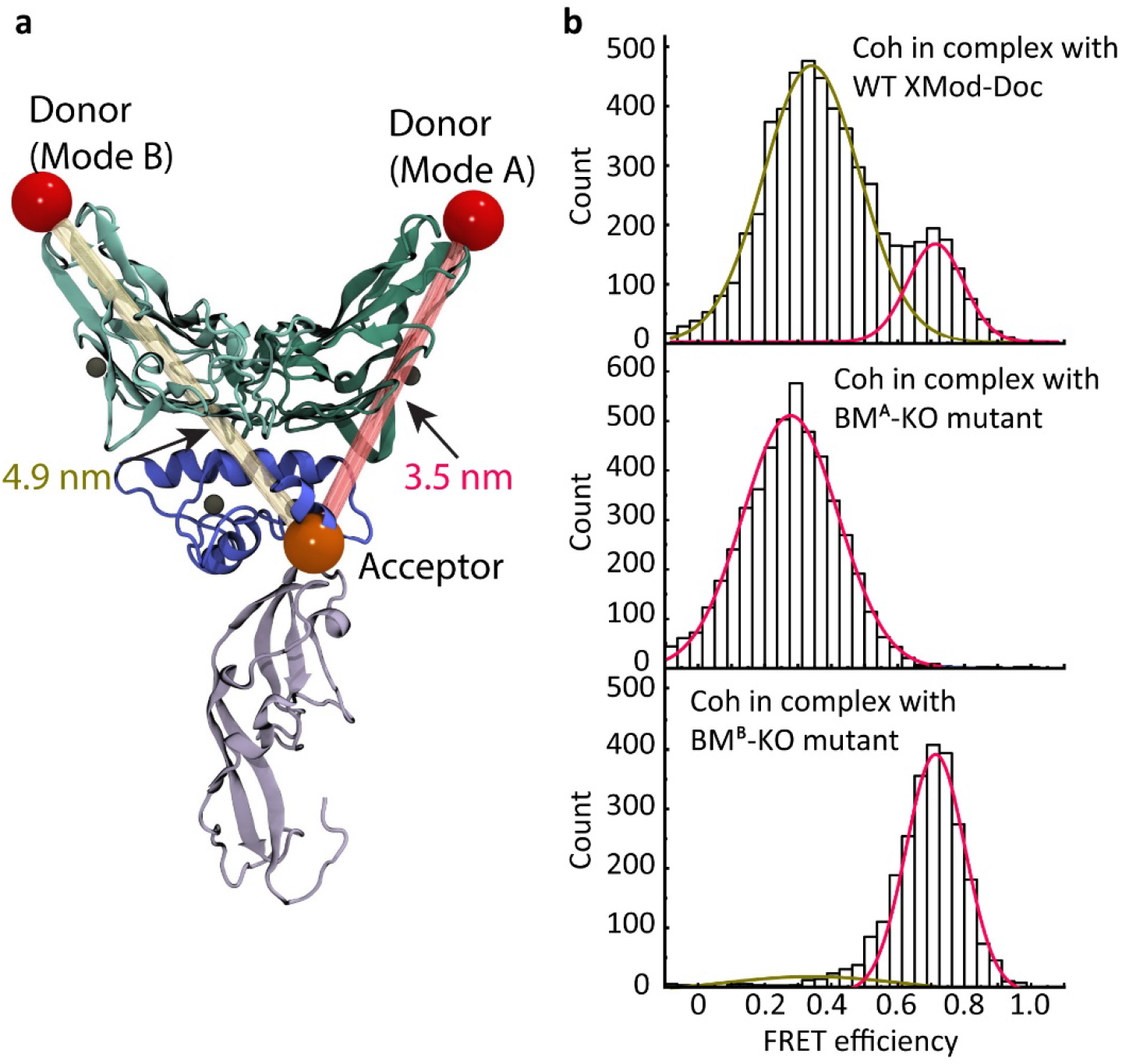
Dual binding modes are observed by smFRET. **a:** The C-terminus of Coh (position 154) was labeled with the FRET donor Cy3b while the C terminus of the XMod-Doc (position 199) was labeled with the FRET acceptor AF647. The donor-acceptor distance was smaller in binding mode A (3.5 nm) than the binding mode B (4.9 nm), giving rise to a higher FRET efficiency when the complex formed in binding mode A. **b:** FRET efficiency histograms measured using Cy3b-labeled WT XMod-Doc (top), BM^A^-KO (middle), or BM^B^-KO (bottom) complexed with AF647-labeled Coh. Efficiency histograms were fitted with single or double Gaussian distributions.

### Kinetic model and Monte Carlo simulations

Combining the experimental results and MD simulations led us to propose a kinetic scheme for the unbinding mechanism of the *Rc* XMod-Doc:Coh complex that accounts for dual-binding modes as well as the catch bond behavior observed under force ramp conditions (Fig. 5a). Our model postulates that there are two non-interconvertible bound states with different mechanical stabilities (binding modes A and B). Upon binding, the complex has an 80% probability of forming the more stable binding mode A, and a 20% probability of forming binding mode B. If the complex forms in binding mode B, the only escape pathway under force load is P3 terminating in a low force rupture (~200 pN). When bound in binding mode A, the complex either ruptures at high force (P1), or enters a weakened state due to the unfolding of XMod (P2). The rate of entering the weaker state (P2) from the stronger state (P1) decreases as the loading rate increases because of the steeper loading rate dependency of XMod unfolding. This results in an increased proportion of P1 high rupture force curves when the complex is probed using force ramp conditions at high loading rates (>100 nN/sec), which is precisely what is observed in classical catch bonds.

**Figure 5.**
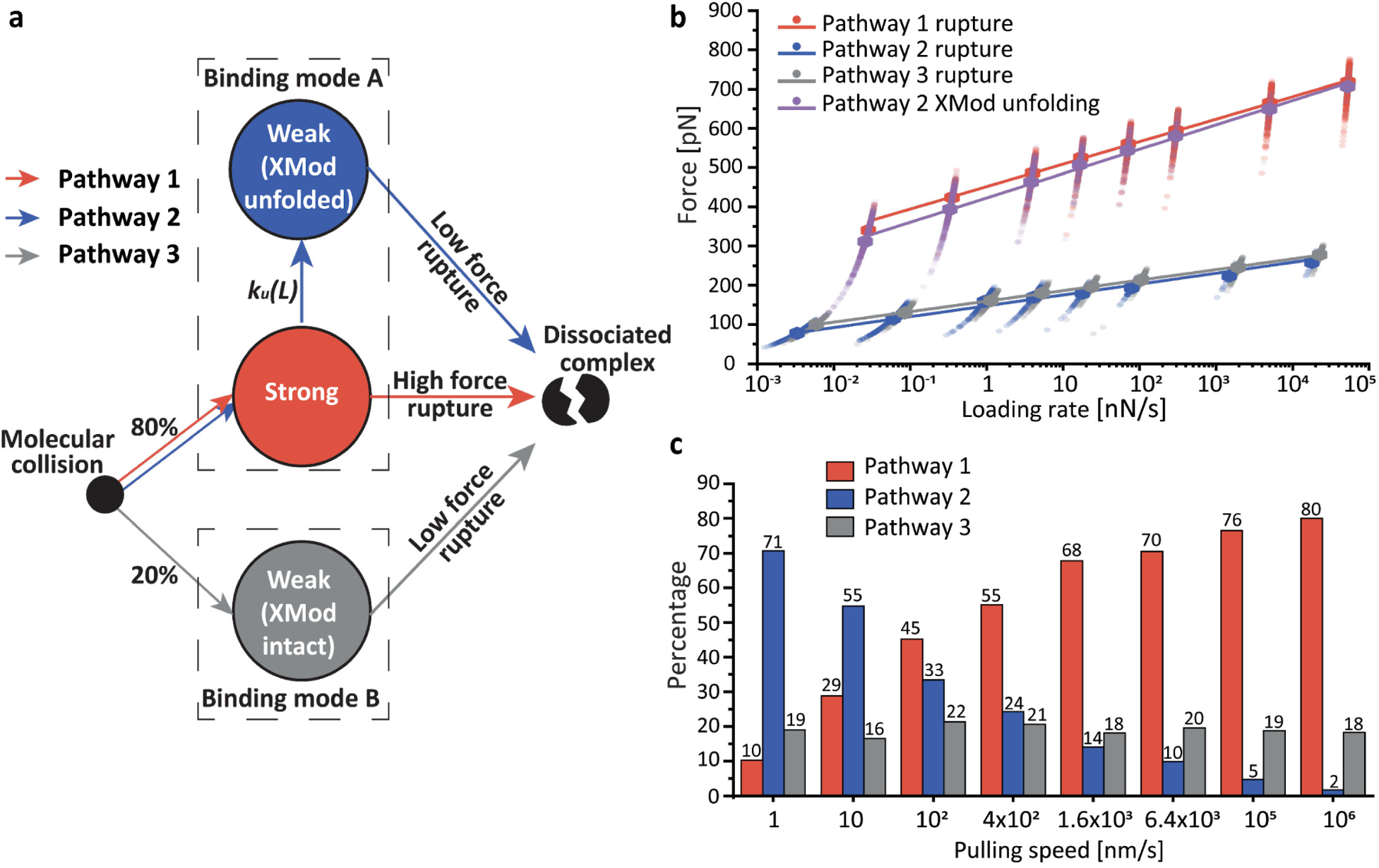
Multi-state kinetic model and Monte-Carlo simulation of mechanical rupture of XMod-Doc:Coh. **a:** The multi-state kinetic model postulates that upon molecular collision, the complex has ~80% probability of forming the strong binding mode A (pathways 1 and 2) and 20% probability of forming the weaker binding mode B (pathway 3). Once the binding mode is set, there is no interconversion between the modes. In the strong binding mode, the complex can rupture at high force (pathway 1) with XMod remaining folded, or XMod can unfold prior to complex rupture according to a loading rate dependent unfolding rate *k*_*u*_*(L)* (pathway 2). In pathway 2, XMod unfolding destabilizes the complex, resulting in low force complex rupture. At increased loading rates, the XMod unfolding rate decreases so that pathway 2 is deactivated and the complex has a higher probability of unbinding along pathway 1 (high force rupture). In pathway 3, the complex ruptures at low force without reaching sufficiently high force to unfold XMod. **b and c:** Monte-Carlo simulation results. Force *vs.* extension curves at constant pulling speed were simulated to obtain the loading rate dependency of complex rupture and XMod unfolding events (**b**), as well as the percentages of the three unbinding pathways at different pulling speeds (**c**). The simulation was carried out at same pulling speeds as the experiments (100, 400, 1600, 6400 nm/s) and further extended over a range from 1 to 10^6^ nm/s. Simulated force *vs.* loading rate plots were fitted with the Bell-Evans model to extract *k*_*0*_ and *Δx*^*‡*^ values (**Table S1**).

We used kinetic rates obtained from AFM-SMFS combined with our proposed state model to simulate the system in a constant pulling speed scenario, identical to the experiments. We used worm-like chain elasticity theory and Monte-Carlo^59^ to simulate force-extension curves of the XMod-Doc:Coh stretching, XMod unfolding and complex rupture (see Supplementary Information). The loading rate dependency of the complex rupture force and XMod unfolding forces, as well as the rupture and unfolding force histograms from the simulated curves are shown in Fig. 5b, **S8** and **S9**. The simulations showed remarkable agreement with experiment results both in terms of the rupture forces, and other observed trends (Fig. 2, **S3** and **S4**). For example, our network Monte-Carlo modeling shows the same bimodality of the rupture force distributions, similar force magnitudes and similar ratios between the P1, P2, and P3 trajectories. Furthermore, the novel catch bond network topology that emerged in force ramp mode was also observed in the simulation.

The simulations further allowed us to probe a range of pulling speeds that were not accessible experimentally. We extended the range of pulling speeds in the simulations to get a clearer picture of the novel catch bond behavior. As shown in Fig. 5c, at high loading rates, the complex predominantly ruptures along P1 due to the strengthening of XMod. At extremely slow pulling speeds, we see in the simulation that the P1 pathway is lost and the complex only exhibits P2 and P3 low force rupture behavior. The broad agreement of the simulation with the experimental results provided strong support for the proposed kinetic scheme.

## Conclusions

We discovered a new mechanism by which bacteria achieve mechanically stable adhesion to crystalline fiber surfaces in the human gut, and resolved the dual binding modes of this complex using single-molecule techniques. The kinetic scheme amounts to a novel multi-state catch bond mechanism in binding mode A (P1/P2 paths). The system starts in the strong (P1, activated) state and has a certain probability of entering the weak state (P2). The transition rate from P1 to P2 decreases with increasing loading rate, meaning that the weak state is inhibited at high loading rates. Once the complex enters the weak state, it cannot return to the strong state. These features make our system distinct from the other two-state catch bond models^4,7,12,13,60^. Interestingly, the catch bond behavior emerges from a network of purely slip bonds/folds and only manifests under a force ramp or constant speed scenario. If this system is probed using constant force clamp conditions, there is no increase in lifetime as the clamping force is increased (**Fig. S10**).

Based on structural modelling and analysis, we predicted that the heterogeneity of unbinding pathways was attributable to two different binding conformations, binding modes A and B. AFM-SMFS and smFRET on mutant XMod-Doc constructs designed to specifically knockout binding mode A or B supported the presence of dual-binding modes with different mechanical properties. The biological significance of the two binding modes is still unclear, however we speculate that the *Rc* bacterium might switch between the low (P3) and high force catch (P1/P2) adhesion modes based on environmental factors or post-translational modifications. Our research demonstrates an entirely new degree of complexity by which bacteria regulate adhesion strength through molecular mechanisms such as dual-binding modes, mechanical allostery, and catch bonding.

## Associated content

Supplementary information (supplementary figures S1-S10, supplementary table S1 and supplementary notes).

## Notes

The authors declare no competing financial interest.

### Acknowledgement

This work was supported by the University of Basel, ETH Zurich, an ERC Starting Grant (MMA-715207), the NCCR in Molecular Systems Engineering, and the Swiss National Science Foundation (Project 200021_175478). A. M. V. has received funding from the European Union’s Horizon 2020 research and innovation program under the Marie Skłodowska-Curie grant agreement No 746635. R.C.B. is supported by the National Institutes of Health (NIH) grant P41-GM104601, and by the National Science Foundation (NSF) grant MCB-1616590. Molecular dynamics simulations made use of GPU-accelerated nodes of NCSA Blue Waters supercomputer as part of the Illinois allocation grant “ILL_baxs”. The state of Illinois and the National Science Foundation (awards OCI-0725070 and ACI-1238993) support Blue Waters sustained-petascale computing project. P. T. acknowledges support by the DFG (excellence cluster CIPSM (Center for Integrated Protein Science Munich), INST 188/401-1 FUGG). The authors thank Peter Schultz for providing the pEVOL-pAzF plasmid, Duy Tien Ta for providing plasmid carrying the elastin-like peptide and Mariia Beliaeva for help with ITC measurement and ITC data analysis. The authors further thank Lukas Milles and Markus Jobst for helpful discussions.

## Materials and methods

All reagents were at least of analytical purity grade and were purchased from Sigma-Aldrich (St. Louis, MO, USA), Thermo Fisher Scientific (Waltham, MA, USA), GE Healthcare (Chicago, IL, USA), New England Biolabs (Ipswich, MA, USA) or ABCR GmbH (Karlsruhe, Germany).

Synthetic genes were purchased from Thermo Fisher Scientific.

Primers for cloning were purchased from Microsynth AG (Balgach, Switzerland).

All buffers were filtered through a 0.2 μm polyethersulfone membrane filter (Sarstedt, Nuembrecht, Germany) prior to use. The pH of all buffers was adjusted at room temperature.

## Homology modeling and molecular dynamics simulations

To the best of our knowledge, the structure of the XMod-Doc domain from Scaffoldin B of *Ruminococcus champanellensis* and its binding partner, the CohE domain from the same bacterium, has not been solved by experimental means. Using homologous structures available in the Protein Data Bank (www.pdb.org), we employed Modeller 9.22^38^ to obtain a homology model of the two *R. champanellensis* cellulosomal domains. Modeller works by setting spatial restriction to the atomic positions of the model protein, based on 3D-template structures. Using the *Rc* XMod-Doc:Coh protein sequences we performed a protein BLAST^61^, finding one satisfactory homologue template for the *Rc* XMod-Doc domain, and two for the *Rc* Coh domain. For all the templates, the sequence identity was observed to be low: 20% identity between *Rc* XMod-Doc and the *R. flavefaciens* XMod-Doc (PDB 4IU3) template; 15% identity between *Rc* Coh domain and the *Rf* Coh E (PDB 4IU3) template; 18% identity between *Rc* Coh domain and the *Rf* Coh G (PDB 4WKZ) template. Likewise, the sequence similarities were also found to be small, with 35%, 28%, and 34% respectively. Regarding the templates, both the 4WKZ^40^ and the 4IUI3^39^ structures were solved by means of x-ray crystallography, with a resolution of 1.79Å and 1.97Å, respectively.

Using Modeller, we generated 10 structural models for the *Rc* XMod-Doc domain based on its template, and 20 structural models for the *Rc* Coh domain based on its two templates (10 models for each template). Using VMD^41^, the structure of the *Rf* XMod-Doc:Coh complex was used as a guide to fit all 200 possible combinations of the *R. champanellensis* model structures into binding mode A. For the binding mode B, first an inverted *Rf* XMod-Doc:Coh binding was created by superimposing Doc helix 1 with helix 3, and helix 3 with helix 1, creating a 180° rotated Coh structure. VMD was then used again to fit all the possible 200 models of *R. champanellensis* to this inverted *R. flavefaciens* structure. Typically the best structural model could be selected by employing tools like PROCHECK^62^ and ERRAT server^63^, however, due to the low sequence identity and similarity, we adopted a strategy of using molecular dynamics (MD) to thoroughly test all the homology models.

Employing QwikMD^50^, all 400 model structures were subjected to 5 ns of equilibrium MD to ensure conformational stability. Although after a visual inspection we could see that many of the structural models were not stable following MD simulation, we chose to use a more systematic metric to select the best structural models, namely, selecting, for each of the binding modes, the 5 most stable models under load. For that we performed 20 ns of steered molecular dynamics (SMD) for each of the 400 model structures, pulling the complex apart. The simulations revealed that the complexes would rupture at a wide-range of forces, and the 5 models with highest rupture forces for each binding mode were selected as the best models.

To investigate the stability of the best structural models, we performed another set of SMD simulations using the 5 best models as initial structures in what we call an *in silico* force-spectroscopy approach^34^. Using a wide-sampling strategy, 200 steered molecular dynamics (SMD) replicas were carried out for a total of 4 µs for each binding mode, using the 5 different initial structures. All SMD simulations^3^ were performed with a constant velocity protocol using 5.0 Å/ns as the pulling speed. In all simulations, SMD was employed by restraining the position the N-terminal of XMod-Doc domain, while pulling on the C-terminus of the Coh domain.

In our study, all MD and SMD simulations were performed employing the NAMD molecular dynamics package^48,49^. The CHARMM36 force field^64^ was employed to describe all simulations, using an explicit TIP3 water model^65^. Simulations were performed at the NpT ensemble, in periodic boundary conditions. Temperature was kept at 300K using Langevin dynamics for temperature coupling, while a Langevin piston was employed to hold pressure at 1 bar. A distance cut-off of 14.0 Å was applied to short-range, non-bonded interactions, whereas the particle-mesh Ewald (PME) method was employed for long-range electrostatic interactions. The equations of motion were integrated using a 2 fs time step for all simulations performed. All simulations were analyzed using VMD^41^ and its plugins. Surface contact areas of interacting residues/domains were studied using PyContact^66^.

## Cloning

The constructs for AFM measurements were ybbr-ELP-ddFLN4-XMod-Doc-HIS and Coh-ddFLN4-ELP-HIS-ybbr. A ddFLN4 domain was inserted into a pET28a vector containing ybbr-HIS-ELP (for XMod-Doc) or ELP-HIS-ybbr (for Coh) so that the ELP linker was located between the ddFLN4 and the ybbr tag. The XMod-Doc synthetic gene was inserted to the C terminus of ddFLN4 using Gibson assembly and the Coh synthetic gene was inserted to the N terminus of ddFLN4 using restriction digestion cloning (NdeI and BamHI sites). The sequences of the inserted genes were confirmed by Sanger sequencing (Microsynth AG). The His-tag on the XMod-Doc constructs were then moved to the C terminus of the construct.

Protein samples for ITC measurement were prepared by removing ELP and ddFLN4 domains from the AFM measurement constructs.

The Coh smFRET construct was prepared by adding an Avi-tag to the N terminus of the ITC construct and introducing E154C mutation to Coh.

The WT Doc smFRET construct was prepared by replacing the serine at position 199 with an Amber codon (TCC→TGA). The smFRET constructs of the Doc binding mode mutants were prepared by adding the same mutations as the corresponding AFM constructs to the WT Doc smFRET construct.

## Protein expression and purification

All protein samples used for AFM and ITC as well as Coh used in smFRET were expressed in NiCo21 (DE3) cells (New England Biolabs). Cells were cultured in TB (terrific broth) medium containing 50 μg/mL kanamycin until OD600 reached ~0.6. Protein expression was induced by adding 0.5 mM IPTG to the culture, followed by incubating at 20 °C overnight. Cells were harvested and lysed using sonication. The cell lysate was pelleted and the supernatant was loaded onto a His-tap FF 5 mL column (GE Healthcare) and washed with TBS buffer supplemented with calcium (TBS-Ca, 25mM Tris, 72mM NaCl, 1 mM CaCl_2_, pH 7.2). Bound protein was eluted using TBS-Ca buffer containing 500 mM imidazole. Eluted protein was further purified using size-exclusion column (GE Healthcare). Protein solutions for long term storage were concentrated using a Vivaspin 6 centrifugal filter (GE Healthcare) and stored in 35 % (v/v) glycerol at −20 °C. The concentration of the protein stocks were determined to be ~40 µM using UV absorption spectrophotometry.

## Amber suppression

The Doc smFRET constructs were expressed in BL21Star (DE3) cells using amber codon suppression^58^. The pET28a vector carrying the Doc smFRET construct was co-transformed with plasmid pEVOL-pAzF (a gift from Peter Schultz, Addgene plasmid # 31186). Cells were grown in LB medium containing 50 μg/mL kanamycin and 25 μg/mL chloramphenicol until OD600 reached ~0.8. Cells were then pelleted, washed with M9 minimal medium and resuspended in M9 medium containing 50 μg/mL kanamycin, 25 μg/mL chloramphenicol, 0.2 mg/ml p-azido-l-phenylalanine (pAzF) and 0.02% arabinose. The culture was incubated at 37 °C for 1 h and then 1 mM IPTG was added to the culture, followed by incubating at 16 °C overnight. The expressed protein was extracted and purified using the same protocol as for the AFM constructs.

## AFM sample preparation

The preparations of AFM measurement samples were conducted according to previously published protocols^52^. Biolever mini AFM cantilevers (Bruker, Billerica, MA, USA) and cover glasses were cleaned by UV-ozone treatment (cantilevers) or piranha solution (cover glasses), and silanized using (3-aminopropyl)-dimethyl-ethoxysilane (APDMES) to introduce amine groups on the surface. The silanized cantilevers and cover glasses were subsequently incubated with 10 mg/mL sulfosuccinimidyl 4-(N-maleimidomethyl)cyclohexane-1-carboxylate (sulfo-SMCC) solution for 30 min at room temperature in order to introduce maleimide groups on the surface. After incubating with sulfo-SMCC, the cantilevers and glasses were cleaned with ddH_2_O and immediately incubated with 20 mM coenzyme A (CoA) solution for 2 h at room temperature and then cleaned with ddH_2_O. CoA-coated cantilevers and cover glasses were incubated with Coh-ddFLN4-ELP-ybbR and ybbR-ELP-ddFLN4-XMod-Doc fusion proteins, respectively, in the presence of ~5 μM Sfp (phosphopantetheinyl transferase) enzyme for 2 h at room temperature. After incubation, cantilevers and glass surfaces were intensively rinsed with TBS-Ca buffer and stored under TBS-Ca before the measurement.

## AFM-SMFS measurements

SMFS measurements were performed on a Force Robot AFM (JPK instruments, Berlin, Germany). Cantilever spring constants (ranging from 0.07 N/m to 0.1 N/m) were calibrated using the contact-free method. A control experiment was done showing that the contact-free calibration method gave the same result as contact-based method. The cantilever was brought into contact with the surface and withdrawn at constant speed ranging from 100 nm/s to 6400 nm/s. In a typical measurement around 5,000-10,000 force-extension curves were obtained with a single cantilever in an experimental run of 10-20 h. The majority of the data were unusable curves due to lack of interactions, multiple interactions or nonspecific adhesion of molecules to the cantilever tip. However, ~10% of the curves showed single-molecule interactions. We filtered the data by searching for the two-step unfolding patterns and the 64 nm contour length increment of two ddFLN4 fingerprint domains.

## AFM data analysis

AFM data were analyzed using a combination of Python scripts, R scripts (R foundation, available at https://www.r-project.org/, utilizing packages readr and ggplot2 and user interface R Studio, available at https://www.rstudio.com/), and Origin 2018 (OriginLab).

Force-extension curves were transformed into contour length space using freely rotating chain (FRC) model, which assumes bonds of length *b* are connected by a fixed angle *ϒ*. The force-extension curves were transformed to contour length *L* using Eq. S1^67^:

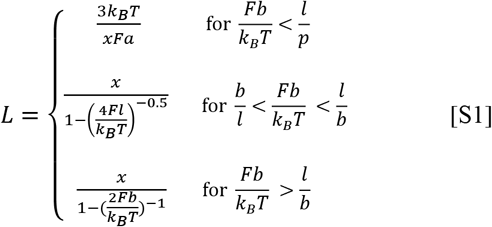

where 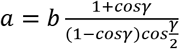 is the Kuhn length and 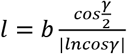 is the persistence length.

The force-extension curves were screened using the ~64 nm contour length increment of two ddFLN4 fingerprint domains.

The most probable rupture force of the complex and unfolding force of XMod was fitted linearly against the logarithm of loading rate to extract the zero-force off rate *k*_*0*_ and the distance to the energy barrier *Δx*^*‡*^ using Eq. S2, as explained by the Bell-Evans model^1,2^:

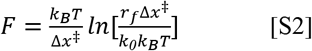

, where k_B_ is the Boltzmann constant and T is the temperature.

The rupture force of the complex and unfolding force of XMod were plotted into histograms and transformed into off rate values using the Dudko-Hummer-Szabo model^56,57^, as explained below.

Histograms were plotted using equal bin width *ΔF* = 40 pN. For one histogram containing N bins, starting from *F*_*0*_ and ending at *F*_*N*_ = *F*_*0*_ + *NΔF*, the k_th_ bin can be directly transformed into the force-dependent rate constant value using Eq. S3:

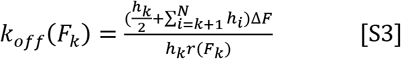

where *k*_*off*_(*F*_*k*_) is the off rate under the average unfolding or rupture force of the k_th_ bin, *r*(*F*_*k*_) is the average loading rate of the k_th_ bin, *h*_*k*_ is the height of the k_th_ bin, which is calculated using Eq. S4:

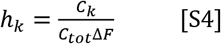

where *C*_*k*_ is the number of counts in the k_th_ bin and *C*_*tot*_ is the total number of counts in the histogram.

Based on Kramers theory, the force-dependence of *k*_*off*_(*F*) can be written as Eq. S5:

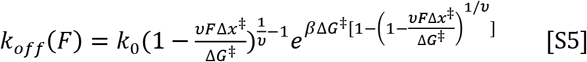

where *k*_*0*_ is the intrinsic off rate in the absence of force, *Δx*^*‡*^ is the distance to the energy barrier, *ΔG*^*‡*^ is the height of the energy barrier in the absence of force, β^−1^=k_B_T, and *υ* = 0.5, which assumes the shape of the free-energy surface is cusp.

### Single-molecule FRET sample and chamber preparation

The Coh smFRET construct was reduced by adding 5 mM TCEP (tris(2-carboxyethyl)phosphine) and incubating at room temperature for 30 min. The reduced protein was mixed with 20-fold excess of maleimide-Cy3b (GE Healthcare) and incubated at room temperature for 1 h followed by incubation at 4 °C overnight. The Doc smFRET constructs incorporated with p-azido-phenylalanine were labeled by mixing with 5-fold excess of DBCO-AF647 (Jena Bioscience, Jena, Germany) and incubating at room temperature for 1 h, followed by incubation at 4 °C overnight. The labeled Coh and Doc constructs were purified using a desalting column followed by size-exclusion column.

smFRET experiments were carried out in Lab Tek chambers (Lab-Tek II chambered coverglass system, Thermo Fisher Scientific). Prior to the measurement, chambers were passivated with 1 mg/ml BSA (PAA Laboratories GmbH, Germany) for at least 1 hour. BSA solution was removed only before the measurement and the chamber was washed twice with PBS and once with measurement buffer (50 mM Tris pH 8, 100 mM NaCl, 1 mM CaCl_2_, 2 mM Trolox/Trolox quinone and 1% glucose).

### Single-molecule FRET measurements

Solution smFRET experiments were performed on a PIE-based^68^ home built confocal microscope based on an Olympus IX-71 inverted microscope. Two pulsed lasers (639 nm, 80 MHz, LDH-D-C-640; 532 nm, 80 MHz, LDH-P-FA-530B, both from PicoQuant GmbH) were altered on the nanosecond timescale by a multichannel picosecond diode laser driver (PDL 828 “Sepia II”, PicoQuant GmbH, Berlin, Germany) with an oscillator module (SOM 828, PicoQuant GmbH). The lasers were coupled into a single mode fiber (P3-488PM-FC, Thorlabs GmbH, Dachau, Germany) to obtain a Gaussian beam profile. Circular polarized light was obtained by a linear polarizer (LPVISE100-A, Thorlabs GmbH) and a quarter-wave plate (AQWP05M-600, Thorlabs GmbH, Dachau, Germany). The light was focused by an oil-immersion objective (UPLSAPO100XO, NA 1.40, Olympus Deutschland GmbH) onto the sample. The sample was moved by a piezo stage (P-517.3CD, Physik Instrumente (PI) GmbH & Co. KG, Karlsruhe, Germany) controlled by a E-727.3CDA piezo controller (Physik Instrumente (PI) GmbH & Co. KG, Karlsruhe, Germany). The emission was separated from the excitation beam by a dichroic beam splitter (z532/633, AHF analysentechnik AG) and focused onto a 50 μm pinhole (Thorlabs GmbH). The emission light was split by a dichroic beam splitter (640DCXR, AHF analysentechnik AG) into a green (Brightline HC582/75, AHF analysentechnik AG; RazorEdge LP 532, Laser 2000 GmbH) and red (Shortpass 750, AHF Analysentechnik AG; RazorEdge LP 647, Laser 2000 GmbH) detection channel. Emission was focused onto avalanche photodiodes (SPCM-AQRH-14-TR, Excelitas Technoligies GmbH & Co. KG) and signals were registered by a time-correlated single photon counting (TCSPC)-unit (HydraHarp400, PicoQuant GmbH, Berlin, Germany). The setup was controlled by a commercial software package (SymPhoTime64, Picoquant GmbH, Berlin, Germany). Excitation powers of 36 µW and 25 µW were used for donor and acceptor lasers (as measured in front of the entrance of the microscope).

Labeled Coh and Doc samples were mixed in a molar ratio of 1:1 at a concentration of 1 µM, incubated for 1 minute, and finally diluted in the chamber to a concentration of 200 pM.

### Single-molecule FRET data analysis

smFRET burst selection was performed using a sliding time window burst search algorithm, with a time window of 500 µs and a minimum of 4 photon per time window. A threshold for burst detection of 40 photons was used^69^. In order to sort out photobleaching and blinking events, ALEX-2CDE^70^ and ׀TDX-TAA׀ filters^71^ were used. Doubled-labeled Doc:Coh complexes were further selected by keeping the stoichiometry parameter between 0.2 and 0.8. Accurate FRET efficiencies^29,72^ were calculated from fluorescence intensities as:

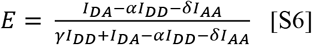

where *I*_*DA*_, *I*_*AA*_ and *I*_*DD*_ are the background-corrected photon counts in the acceptor channel after donor excitation, the acceptor channel after acceptor excitation, and the donor channel after donor excitation. The *α* and *δ* correction parameters are calculated from donor only and acceptor only subpopulations and accounts for spectral cross talk and direct excitation of the donor dye. The different detection efficiencies and quantum yields of fluorophores are corrected with the γ correction factor^29,72^.

### ITC measurement

The titration was carried out at 25 °C using a VP-ITC instrument^73^. The analyte was 16.1 µM Coh (lacking ELP linker and ddFLN4 domains) and the injectant was 126 µM XMod-Doc protein (lacking ELP linker and ddFLN4 domains). Both protein samples were in TBS-Ca buffer. The titration was carried out by injecting XMod-Doc solution dropwise into the analyte. Each drop contained 10 µL XMod-Doc solution and there was 5 min retention time between two consecutive drops so that the system could equilibrate after injecting a drop. The power required to maintain equal temperature between the sample cell and the reference cell (filled with water) was recorded. The titration was terminated after 27 injections, when the analyte (Coh) was fully saturated by the injectant (Doc).

### Monte Carlo simulation

A Monte Carlo approach based on Kramers theory was used to validate the multi-state kinetic model. The receptor-ligand dissociation in combination with fingerprint domain unfolding was simulated in a constant pulling speed protocol. Briefly, the XMod-Doc:Coh complex was randomly assigned a binding mode to be either binding mode A (80% possibility) or binding mode B (20% possibility). The corresponding kinetic parameters (*k*_*0*_ and *Δx*^*‡*^, see Table 1) extracted from AFM-SMFS were used for the simulation. A series of force values *F(t*_*i*_) was generated on an evenly distributed extension axis *X(t*_*i*_) using a worm-like chain (WLC) model^74^. Due to the fact that the constant pulling speed protocol is achieved by the constant speed pulling of the AFM head instead of the AFM tip, a bending correction was done by converting the molecular extension *X(t*_*i*_) to the AFM head height *H(t*_*i*_) using Eq. S7:

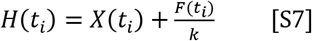

where *k* is the spring constant of the AFM cantilever. Then the time series could be generated based on the pulling speed *V*:

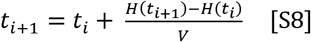

During each time slice(∆*t* = *t*_*i*+1_ − *t*_*i*_), the probability of XMod-Doc:Coh rupture or protein domain unfolding was calculated using the following equation:

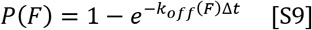

where *k*_*off*_*(F)* can be drawn from Eq. S9 following the Bell-Evans model:

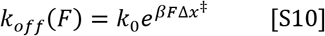

where β = (k_B_T)^−1^. The dissociation probability is compared to a random number between zero and unity. If the random number is smaller than *P(F)* the rupture or unfolding event occurs and the corresponding force is recorded as the rupture or unfolding force. For each pulling speed, 1000 curves were generated and a histogram was drawn for the complex rupture force as well as the XMod unfolding force (**Fig. S8 and S9**). For simulation under force clamp conditions, a constant force was used and 1000 curves were generated to calculate the lifetime of the complex under each applied force (**Fig. S10**).

