## Supplementary Information for "High Force Catch Bond Mechanism of Bacterial Adhesion in the Human Gut"

### 1. Supplementary figures S1-S9

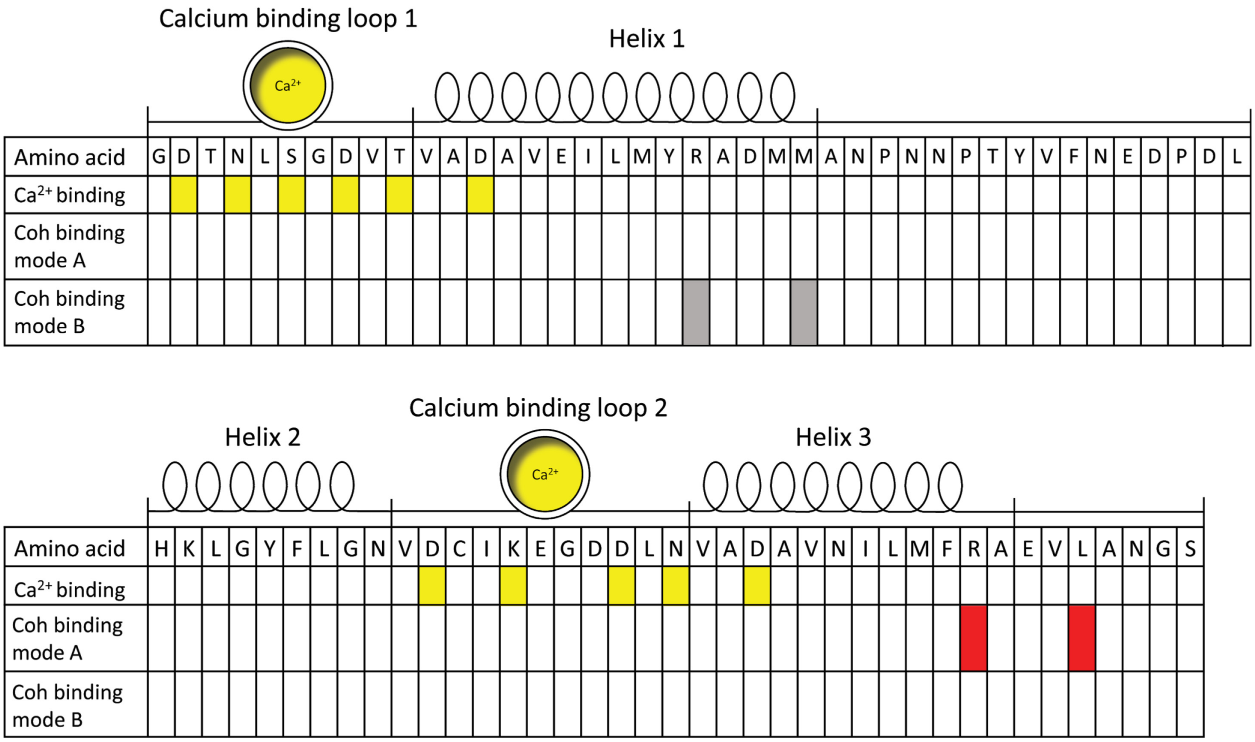

**Figure S1. Amino acid sequence and secondary structure elements of XMod-Doc.**

The residues involved in calcium binding are shown in yellow, the residues mutated to knock out the binding mode A are shown in red and the residues mutated to knock out the binding mode B are shown in grey.

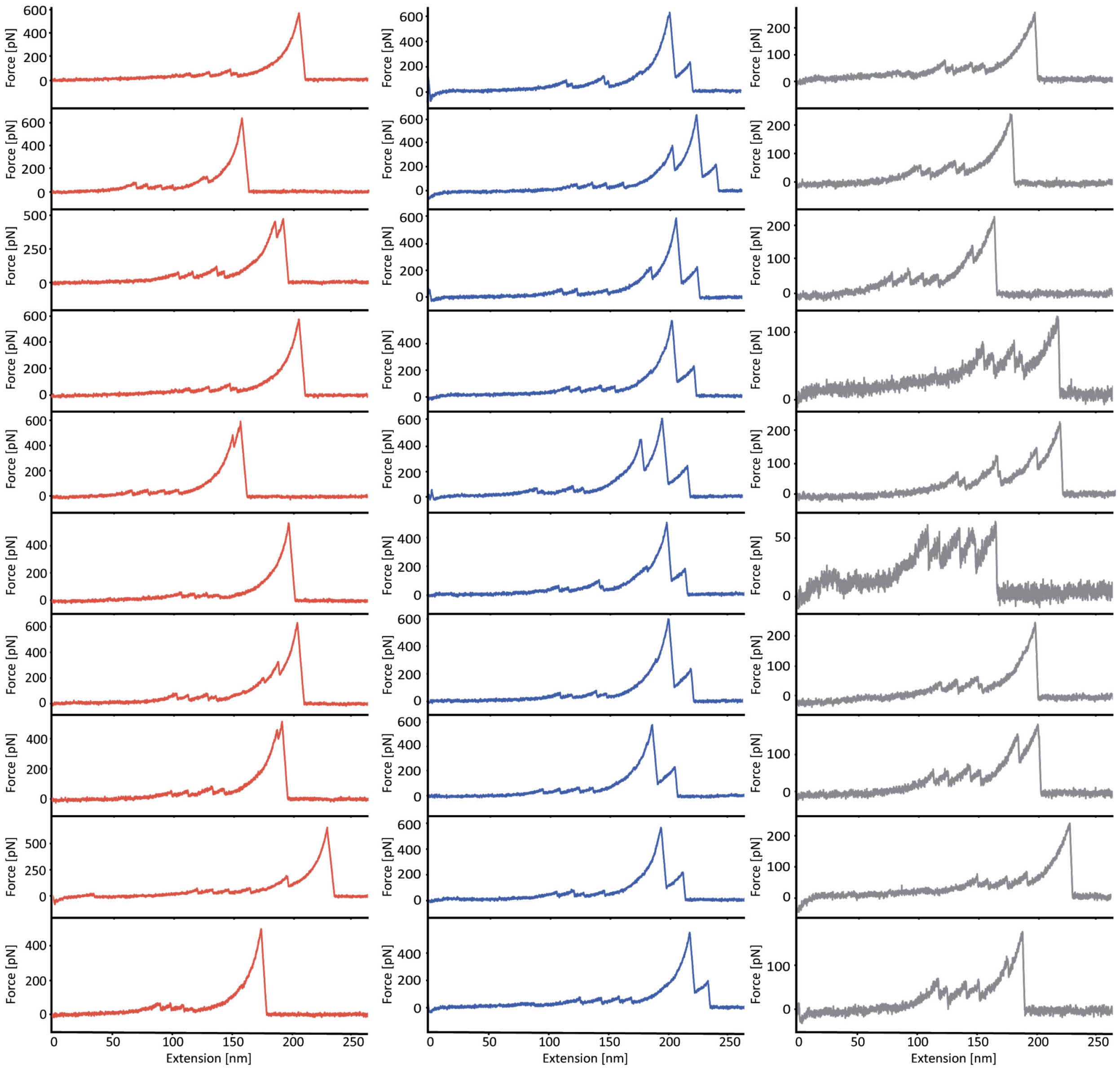

**Figure S2. Example force-extension curves obtained in AFM-SMFS measurements.**

Examples of pathways 1 (left column, red), 2 (middle column, blue) and 3 (right column, grey) force-extension curves obtained in AFM-SMFS measurements of WT XMod-Doc:Coh complex. Some force curves showed unassigned unfolding events between ddFLN4 unfolding and complex rupture or XMod unfolding. These unfolding events broadened the contour length histogram and were attributed to partial unfolding of Coh or Doc domains.

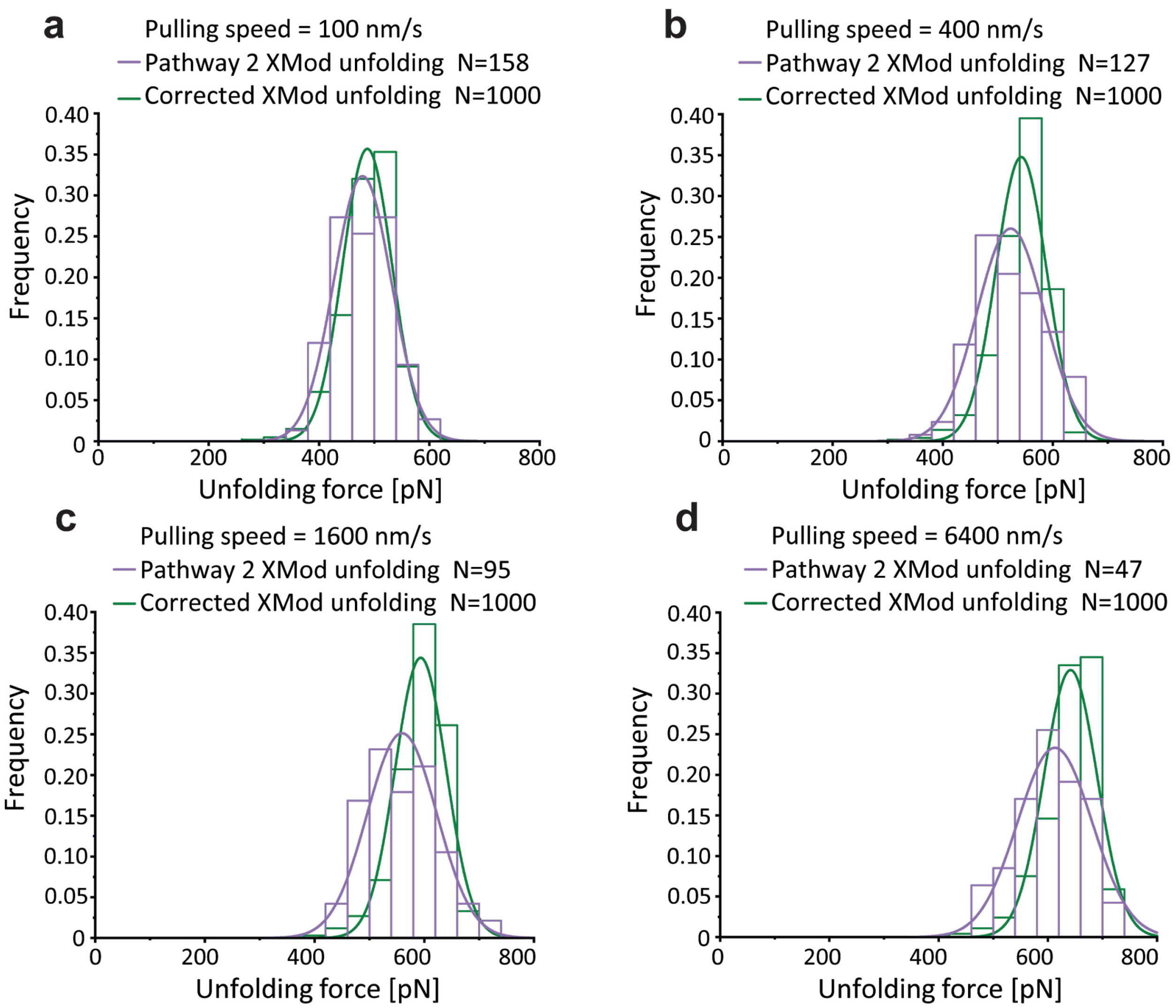

**Figure S3. Correction of XMod unfolding force taking biasing effect into account.**

The XMod unfolding forces measured in pathway 2 under four different pulling speeds were plotted as histograms and fitted with Gaussian distributions (purple). The XMod unfolding force distributions were corrected using iterative fitting which takes the biasing effect[1] into account (green).

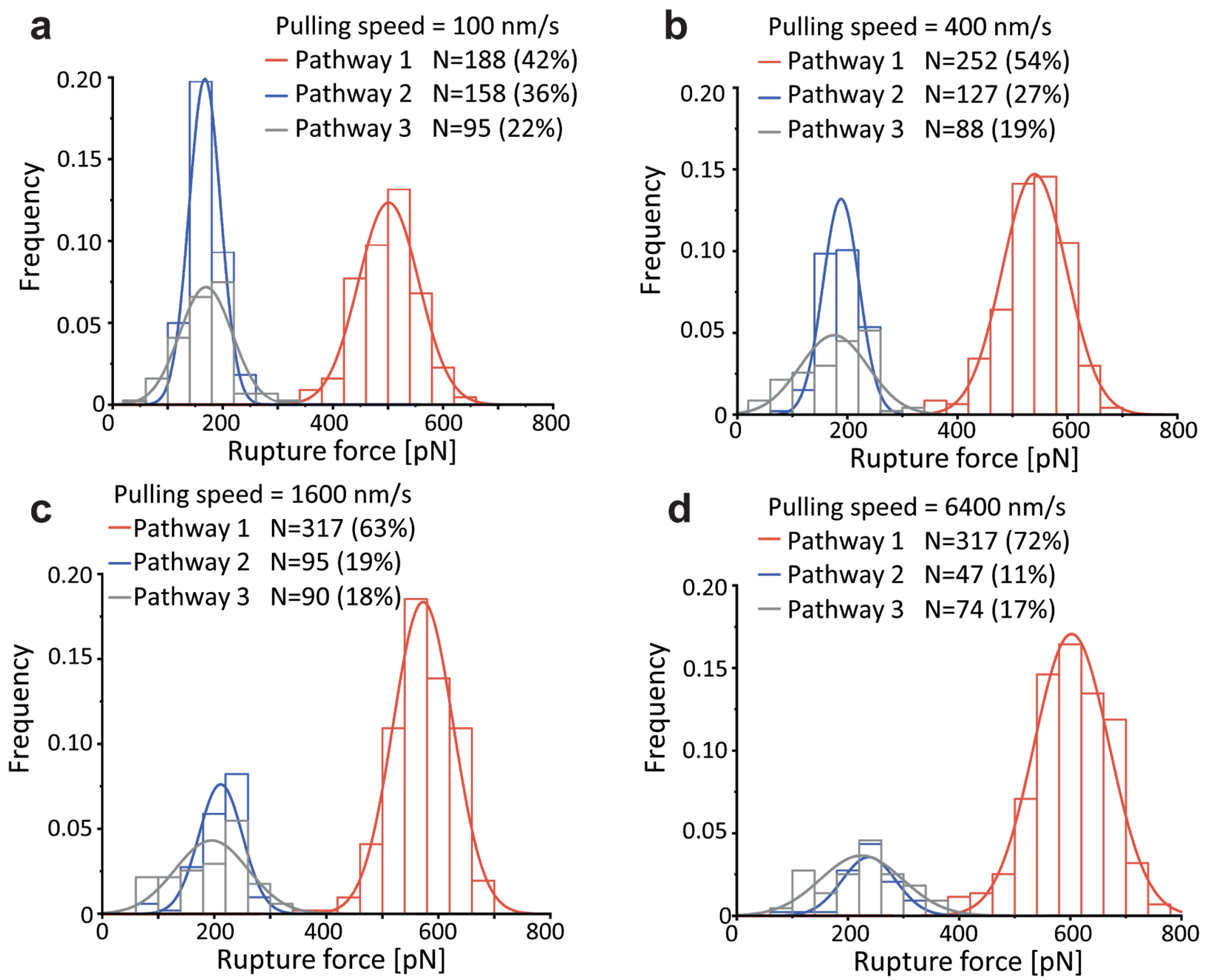

**Figure S4. XMod-Doc:Coh complex rupture force histograms at different pulling speeds.**

The three different pathways are plotted in different colors: High rupture force (P1, red); XMod unfolded, low rupture force (P2, blue); and XMod folded, low rupture force (P3, grey). Each histogram was fitted with a Gaussian distribution. Loss of the P2 population at higher pulling speeds drives catch bond behavior in the force ramp scenario.

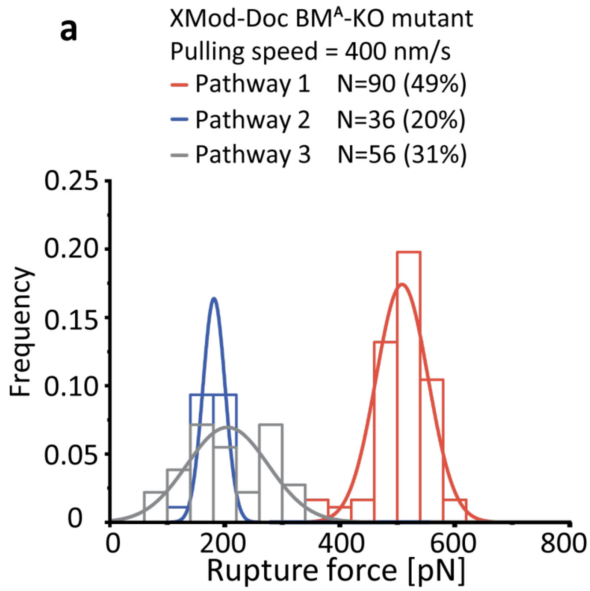

**Figure S5. Rupture force histograms of binding mode mutants.**

**a:** The rupture force histogram measured using BM$^{A}$-KO mutant, showing a decreased percentage of pathways 1 and 2 curves. **b:** The rupture force histogram measured using BM$^{B}$-KO mutant, showing a decreased percentage (nearly total loss) of pathway 3 curves.

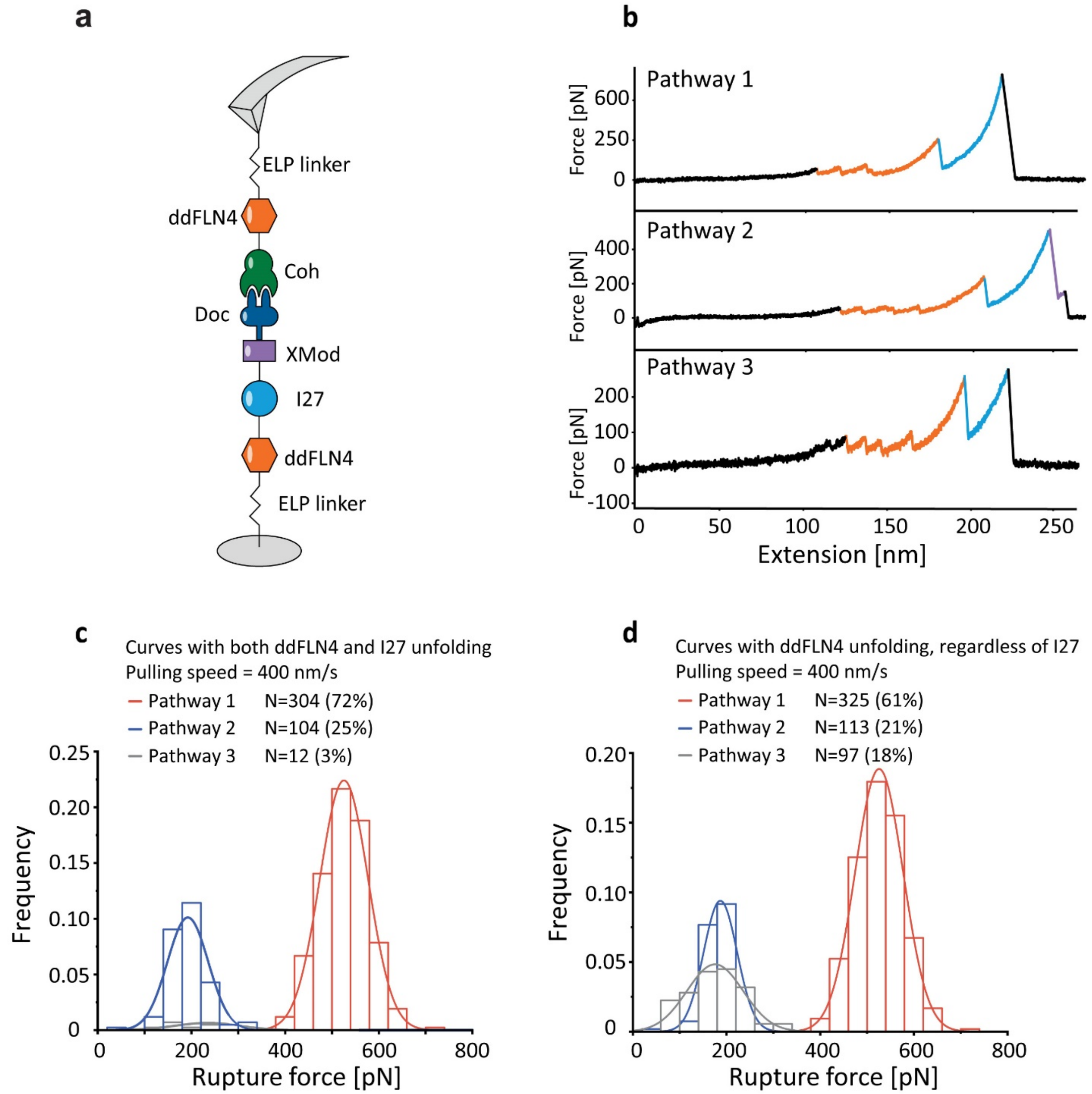

**Figure S6. AFM measurements with I27 fingerprint domain.**

**a:** AFM setup using the I27 biasing effect to demonstrate two binding modes. **b:** Example force-extension curves showing unfolding of 2x ddFLN4 (in orange) and I27 (in blue) in all three pathways. **c:** Rupture force histogram of force curves filtered with both ddFLN4 and I27 fingerprint domains shows that complexes capable of unfolding I27 rarely (3%) dissociated along pathway 3. **d:** Rupture force histogram of force curves filtered with only ddFLN4 showed that pathway 3 was prevalent in the dataset to the same degree as for WT (~18%), but these curves lacked I27 unfolding events.

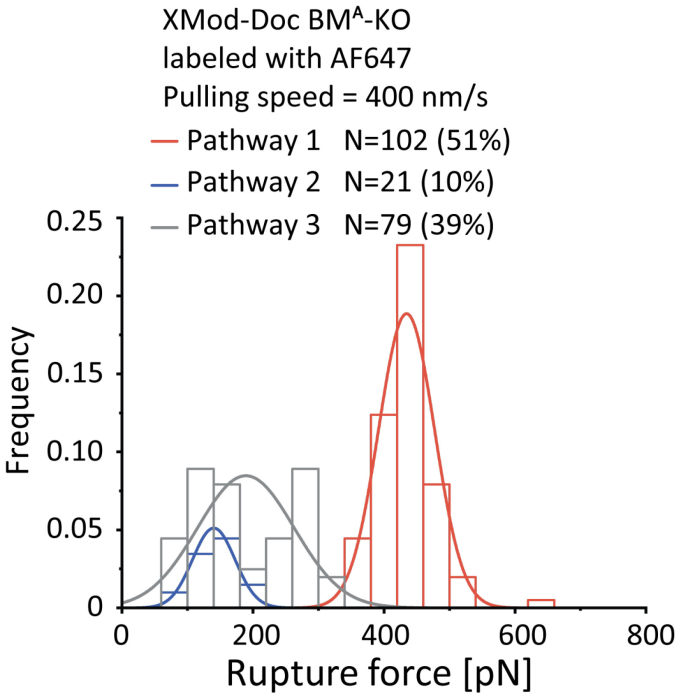

**Figure S7. AFM measurement of AF647 labeled BM$^A$-KO.**

Given the significant molecular weight of the FRET acceptor dye DBCO-AF647 (~1100 grams/mol), we sought to further understand the influence of dye labeling on binding. We used AFM-SMFS to measure the rupture forces between unlabeled Coh and BM$^A$-KO labeled with AF647 at 400 nm/s pulling speed. The most probable rupture force of P1 was found to be 434 pN for the fluorophore labeled BM$^A$-KO complex, which is significantly lower than the P1 rupture force measured using unlabeled BM$^A$-KO (508 pN). We concluded that the AF647 fluorophore at the C-terminus of XMod-Doc slightly destabilized the complex in binding mode A and increased the off-rate in this binding mode. However, fluorophore labeling did not have a significant influence on the rupture force of P3, which corresponds to binding mode B. As a consequence, the binding mode A population was less prevalent than binding mode B in the smFRET measurement. The observed ratios of the two binding modes probed by smFRET reflect the equilibrium scenario, whereas binding mode ratios probed by AFM are governed by differences in on-rates only.

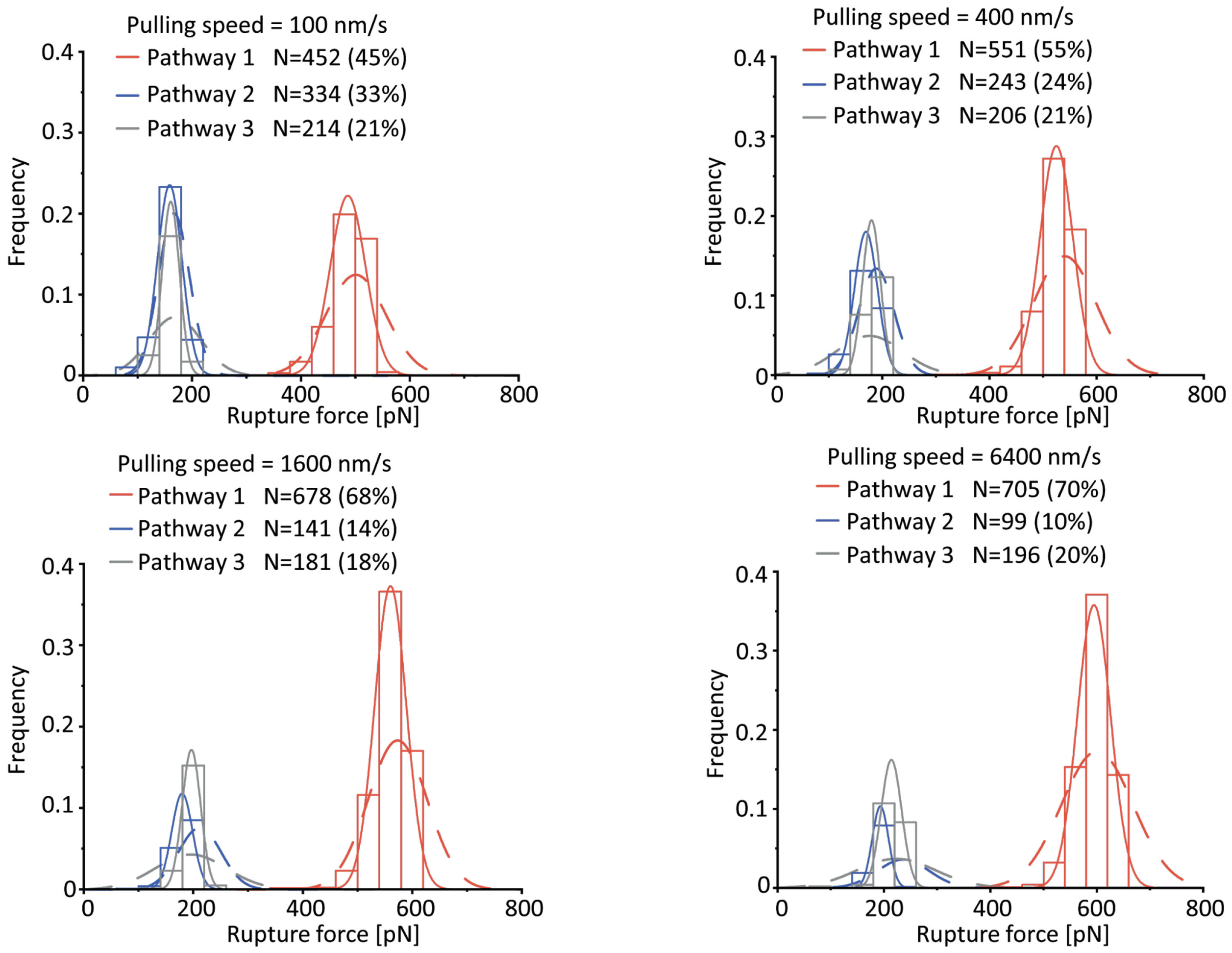

**Figure S8. Monte-Carlo simulation results of complex rupture forces.**

The complex rupture forces were simulated at different pulling speeds and plotted as histograms. The histograms were fitted with Gaussian distributions (solid lines). The corresponding experimental rupture force distributions are shown as dashed lines.

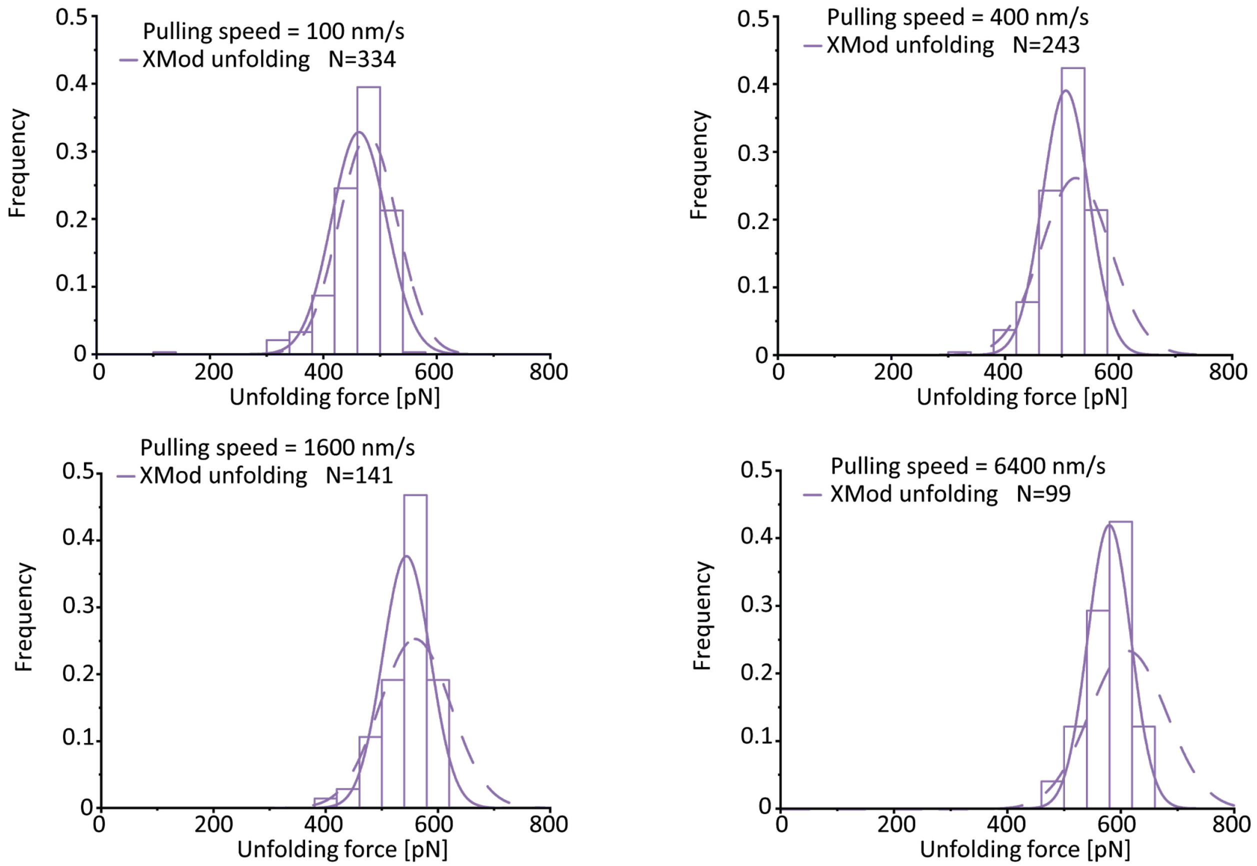

**Figure S9. Monte-Carlo simulation results on XMod unfolding force.**

The XMod unfolding forces in pathway 2 were extracted from Monte Carlo simulations at different pulling speeds and plotted into histograms. The histograms were fitted with Gaussian distributions (solid lines). The corresponding experimental unfolding force distribution is shown as a dashed line.

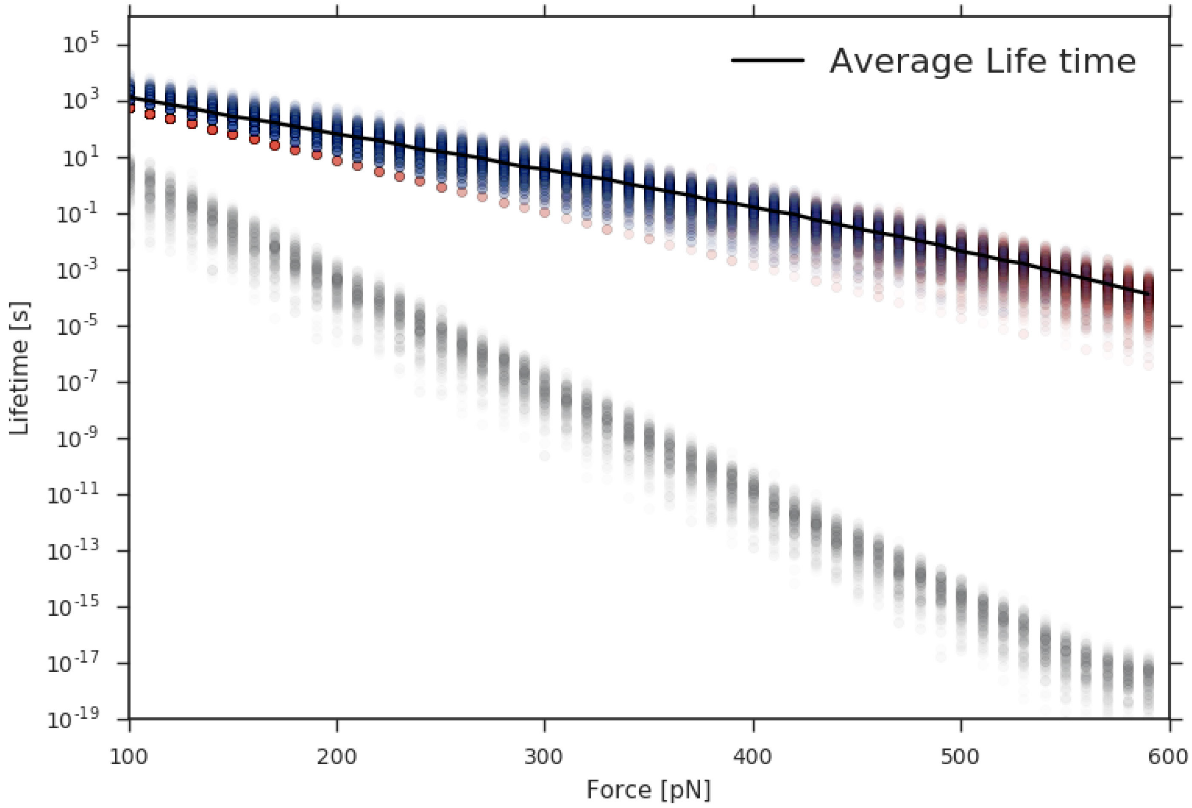

**Figure S10. Force clamp Monte-Carlo simulation.**

The force-dependent lifetime of the complex was simulated in force clamp mode (red: pathway 1, blue: pathway 2, grey: pathway 3). The average lifetime (black line) does not exhibit catch bond behavior for the system probed in force clamp mode.

2. **Supplementary table**

**Table S1. Kinetic parameters extracted from Monte-Carlo simulations**

| Pathway | Event | $k_0$ [$s^{-1}$] (Bell-Evans) | $\Delta x^{\ddagger}$ (Bell-Evans) |
|---|---|---|---|
| 1 | High force rupture (XMod folded) | $5.25 \times 10^{-7}$ | 0.167 |
| 2 | Unfolding of XMod (measured) | $5.85 \times 10^{-6}$ | 0.154 |
| | Low force rupture after XMod unfold | $4.04 \times 10^{-4}$ | 0.344 |
| 3 | Low force rupture (XMod intact) | $9.84 \times 10^{-5}$ | 0.356 |

3. **Supplementary notes**

#### 3.1 Amino acid sequences

Color codes: ybbr-tag, his-tag, ddFLN4, I27, ELP linker, XMod, Doc, Coh, Avi-tag, inter-domain linker, mutations (AzF = p-azido-phenylalanine).

**ybbr-ELP-ddFLN4-XMod-Doc-HIS (wild type)**

GTDSLEFIASKLAWGSGHGVGVPGMGVPGVGVPGVGVPGVGVPGVGVPGVGVPGVGV
PGVGVPGEGVPGEGVPGVGVPGMGVPGVGVPGVGVPGVGVPGVGVPGVGVPGVGVP
GVGVPGEGVPGEGVPGVGVPGMGVPGVGVPGVGVPGVGVPGVGVPGVGVPGVGVPG
VGVPGEGVPGEGVPGWPSGSADPEKSYAEGPGLDGGECFQPSKFKIHAVDPDGVHRTD
GGDGFVVTIEGPAPVDPVMVDNGDGTYDVEFEPKEAGDYVINLTLDGDNVNGFPKTVT
VKPAPGSGSGSGSSGSETTTETSVSQSFSFTEPTRVCYWSHDNRPFDLEGLKVILQETIFII
HEDGTVVEAEEQPAAVDITEQCMTIFCNDKQVATPEDTYETQIYKYMLEFAYEGNPDVI
VGTHQVLIGVKGDTNLSGDVTVADAVEILMYRADMMANPNNPTYVFNEDPDLHKLGY
FLGNVDCIKEGDDLNVADAVNILMFRAEVLANGSATWEQIVGYDLIDPSDESTFRTIEKL
GSHHHHHH

**ybbr-HIS -ELP-ddFLN4-I27-XMod-Doc (wild type)**

GTDSLEFIASKLAHHHHHHWGSGHGVGVPGMGVPGVGVPGVGVPGVGVPGVGVPGV
GVPGVGVPGVGVPGEGVPGEGVPGVGVPGMGVPGVGVPGVGVPGVGVPGVGVPGVG
VPGVGVPGVGVPGEGVPGEGVPGVGVPGMGVPGVGVPGVGVPGVGVPGVGVPGVGV
PGVGVPGVGVPGEGVPGEGVPGWPSGSADPEKSYAEGPGLDGGECFQPSKFKIHAVDP
DGVHRTDGGDGFVVTIEGPAPVDPVMVDNGDGTYDVEFEPKEAGDYVINLTLDGDNV
NGFPKTVTVKPAPLIEVEKPLYGVEVFVGETAHFEIELSEPDVHGQWKLKGQPLAASPD
CEIIEDGKKHILILHNCQLGMTGEVSFQAANTKSAANLKVKELGSGSGSGSSGSETTTETS
VSQSFSFTEPTRVCYWSHDNRPFDLEGLKVILQETIFIIHEDGTVVEAEEQPAAVDITEQC
MTIFCNDKQVATPEDTYETQIYKYMLEFAYEGNPDVIVGTHQVLIGVKGDTNLSGDVTV
ADAVEILMYRADMMANPNNPTYVFNEDPDLHKLGYFLGNVDCIKEGDDLNVADAVNI
LMFRAEVLANGSATWEQIVGYDLIDPSDESTFRTIEKL

**ybbr -ELP-ddFLN4-XMod-Doc-HIS (BM$^{A}$-KO)**

GTDSLEFIASKLAWGSGHGVGVPGMGVPGVGVPGVGVPGVGVPGVGVPGVGVPGVGV
PGVGVPGEGVPGEGVPGVGVPGMGVPGVGVPGVGVPGVGVPGVGVPGVGVPGVGVP
GVGVPGEGVPGEGVPGVGVPGMGVPGVGVPGVGVPGVGVPGVGVPGVGVPGVGVPG
VGVPGEGVPGEGVPGWPSGSADPEKSYAEGPGLDGGECFQPSKFKIHAVDPDGVHRTD
GGDGFVVTIEGPAPVDPVMVDNGDGTYDVEFEPKEAGDYVINLTLDGDNVNGFPKTVT
VKPAPGSGSGSGSSGSETTTETSVSQSFSFTEPTRVCYWSHDNRPFDLEGLKVILQETIFII
HEDGTVVEAEEQPAAVDITEQCMTIFCNDKQVATPEDTYETQIYKYMLEFAYEGNPDVI
VGTHQVLIGVKGDTNLSGDVTVADAVEILMYRADMMANPNNPTYVFNEDPDLHKLGY

FLGNVDCIKEGDDLNVADAVNILMFAAEVEANGSATWEQIVGYDLIDPSDESTFRTIEKL
LGSHHHHHH

**ybbr -ELP-ddFLN4-XMod-Doc-HIS (BM$^{B}$-KO)**

GTDSLEFIASKLAWGSGHGVGVPGMGVPGVGVPGVGVPGVGVPGVGVPGVGVPGVGV
PGVGVPGEGVPGEGVPGVGVPGMGVPGVGVPGVGVPGVGVPGVGVPGVGVPGVGVP
GVGVPGEGVPGEGVPGVGVPGMGVPGVGVPGVGVPGVGVPGVGVPGVGVPGVGVPG
VGVPGEGVPGEGVPGWPSGSADPEKSYAEGPGLDGGECFQPSKFKIHAVDPDGVHRTD
GGDGFVVTIEGPAPVDPVMVDNGDGTYDVEFEPKEAGDYVINLTLDGDNVNGFPKTVT
VKPAPGSGSGSGSSGSETTTETSVSQSFSFTEPTRVCYWSHDNRPFDLEGLKVILQETIFII
HEDGTVVEAEEQPAAVDITEQCMTIFCNDKQVATPEDTYETQIYKYMLEFAYEGNPDVI
VGTHQVLIGVKGDTNLSGDVTVADAVEILMYAADMEANPNNPTYVFNEDPDLHKLGY
FLGNVDCIKEGDDLNVADAVNILMFRAEVLANGSATWEQIVGYDLIDPSDESTFRTIEKL
GSHHHHHH

**ybbr-XMod-Doc (WT, S199AzF)-HIS**

GTDSLEFIASKLAGSSGSETTTETSVSQSFSFTEPTRVCYWSHDNRPFDLEGLKVILQETIF
IIHEDGTVVEAEEQPAAVDITEQCMTIFCNDKQVATPEDTYETQIYKYMLEFAYEGNPDV
IVGTHQVLIGVKGDTNLSGDVTVADAVEILMYRADMMANPNNPTYVFNEDPDLHKLG
YFLGNVDCIKEGDDLNVADAVNILMFRAEVLANGAzFATWEQIVGYDLIDPSDESTFRTI
EKLGSHHHHHH

**ybbr-XMod-Doc (BM$^{A}$-KO, S199AzF)-HIS**

GTDSLEFIASKLAGSSGSETTTETSVSQSFSFTEPTRVCYWSHDNRPFDLEGLKVILQETIF
IIHEDGTVVEAEEQPAAVDITEQCMTIFCNDKQVATPEDTYETQIYKYMLEFAYEGNPDV
IVGTHQVLIGVKGDTNLSGDVTVADAVEILMYRADMMANPNNPTYVFNEDPDLHKLG
YFLGNVDCIKEGDDLNVADAVNILMFAAEVEANGAzFATWEQIVGYDLIDPSDESTFRTI
EKLLGSHHHHHH

**ybbr-XMod-Doc (BM$^{B}$-KO, S199AzF)-HIS**

GTDSLEFIASKLAGSSGSETTTETSVSQSFSFTEPTRVCYWSHDNRPFDLEGLKVILQETIF
IIHEDGTVVEAEEQPAAVDITEQCMTIFCNDKQVATPEDTYETQIYKYMLEFAYEGNPDV
IVGTHQVLIGVKGDTNLSGDVTVADAVEILMYRADMMANPNNPTYVFNEDPDLHKLG
YFLGNVDCIKEGDDLNVADAVNILMFRAEVLANGAzFATWEQIVGYDLIDPSDESTFRTI
EKLGSHHHHHH

**Coh-ddFLN4-ELP-HIS-ybbr**

ADGAAKLSMDQKFAEPGETVEIALNLENFDASWTGLEFLVNYDPKLEVALDGAGDIDY
SYGDAIGAMGKKISVGGAISKDLTADGLKGFAFAWGTATAISGNGQLGVFKFTVPADA
QPGDEFPVNLTVNVGSFIDANKENIPFETVNGWIKIKEEGSGSGSGSADPEKSYAEGPGL
DGGECFQPSKFKIHAVDPDGVHRTDGGDGFVVTIEGPAPVDPVMVDNGDGTYDVEFEP
KEAGDYVINLTLDGDNVNGFPKTVTVKPAPGSGSGSHGVGVPGMGVPGVGVPGVGVP
GVGVPGVGVPGVGVPGVGVPGVGVPGEGVPGEGVPGVGVPGMGVPGVGVPGVGVPG
VGVPGVGVPGVGVPGVGVPGVGVPGEGVPGEGVPGVGVPGMGVPGVGVPGVGVPGV
GVPGVGVPGVGVPGVGVPGVGVPGEGVPGEGVPGWRGHHHHHHGSDSLEFIASKLA

**Avi-Coh (E154C)-HIS**

GLNDIFEAQKIEWHEGSGSADGAAKLSMDQKFAEPGETVEIALNLENFDASWTGLEFLV
NYDPKLEVALDGAGDIDYSYGDAIGAMGKKISVGGAISKDLTADGLKGFAFAWGTATA
ISGNGQLGVFKFTVPADAQPGDEFPVNLTVNVGSFIDANKENIPFETVNGWIKIKCEGSH
HHHHH

#### 3.2 Quantifying dual-binding mode behavior using fingerprint domain biasing effect

As shown in **Fig. S6a**, a titin I27 domain, which under our conditions has an unfolding force around ~200 pN[2,3], was inserted between ddFLN4 and WT XMod-Doc as an additional fingerprint domain. The interaction between XMod-Doc and Coh was then probed using AFM-SMFS in the presence of two ddFLN4 and one I27 fingerprint domain. Unfolding of I27 was identified by its unfolding force of ~200 pN and the contour length increment of ~28 nm, as shown in **Fig. S6b**. Force-extension curves were screened based on the contour length increments given by two ddFLN4 domains and one I27 domain and then sorted into the aforementioned three pathways based on the rupture force of the complex and the folding state of XMod. The rupture force of each pathway was plotted in a rupture force histogram (**Fig. S6c**). In binding mode A, the complex was able to resist an external force up to ~500 pN prior to rupture, which was larger than the force required to unfold I27. Therefore, pathways 1 and 2 were not biased by the additional I27 domain and were still observable in the dataset. However, binding mode B has relatively low mechanical stability and ruptures prior to I27 unfolding. Therefore, the frequency of pathway 3 was significantly decreased to only 3% when using I27 as an additional fingerprint domain for curve selection. However, when screening the force curves only based on the two ddFLN4 domains regardless of whether the curves contained I27 unfolding or not, the frequency of different pathways in the screened curves (**Fig. S6d**) was the same as the construct lacking I27 (**Fig. S4b**). This observation further demonstrated that pathways 1 and 2 (high force curves) belong to a different discrete binding mode than pathway 3 (low force curves). The two binding modes have different mechanical stabilities and cannot be converted to the other one on the timescale of the AFM-SMFS curve (~1 second).
